# Social environment induces reversible remodeling of the adult primate brain

**DOI:** 10.64898/2026.11.19.767871

**Authors:** Daniel J. Gale, Joseph Y. Nashed, Colleen Pearce, Ali Rezaei, Seetha Lakshmi Rajeev, Maryam Ansari Esfeh, Corson Areshenkoff, Ian Goodall-Halliwell, Don Brien, Jonny Smallwood, Valerie A. Kuhlmeier, Mark Sabbagh, Jeffrey D. Wammes, Emily R. Oby, Prameet M. Sheth, Gunnar Blohm, Martin Paré, Stephen Scott, Douglas P. Munoz, Fernanda G. De Felice, Douglas J. Cook, Anita Tusche, Susan Boehnke, Andrew Winterborn, Jason P. Gallivan

**Affiliations:** Centre for Neuroscience Studies, Queen’s University, Kingston, Ontario, Canada; Department of Psychology, Queen’s University, Kingston, Ontario, Canada; McConnell Brain Imaging Centre (BIC), Montreal Neurological Institute, McGill University, Montreal, Quebec, Canada; Department of Biomedical and Molecular Sciences, Queen’s University, Kingston, Ontario, Canada; Department of Pathology and Molecular Medicine, Queen’s University, Kingston, Ontario, Canada; Department of Medicine, Queen’s University, Kingston, Ontario, Canada; Department of Neurosurgery, Queen’s University, Kingston, Ontario, Canada

**Keywords:** Social cognition, social neuroscience, macaques, brain structure, brain connectivity, manifold, longitudinal MRI, functional connectivity

## Abstract

Patterns of social contact can change abruptly, but whether their effects on brain architecture persist or recede when original conditions return remains unclear. Here we followed 13 adult male macaques through three successive phases of single housing, protected-contact paired housing, and restored single housing, acquiring structural and functional MRI and measuring hair cortisol at approximately three-month intervals, with two assessments per phase. Paired housing was associated with grey-matter expansion across temporal association cortex and several subcortical and brainstem structures. Concurrently, default-mode- and limbic-associated regions shifted within a functional-connectivity manifold, while hair cortisol increased. When single housing was restored, grey-matter and cortisol elevations receded, and regional functional changes reversed direction. These trajectories identify a reversible component of adult primate brain architecture that tracks current social conditions.

## INTRODUCTION

Across primates, social relationships are powerful predictors of health, longevity, and reproductive success. Yet these relationships unfold in changing environments, where opportunities for social contact can expand, contract, or be abruptly disrupted. In humans, weak social ties and isolation are associated with poor health and elevated mortality [1–4]. In nonhuman primates, strong social bonds predict infant survival and reproductive success, as well as buffer physiological responses to stress [5–7]. Navigating and maintaining these consequential relationships engages a distributed “social brain” network spanning temporal association cortex, medial and dorsolateral prefrontal cortex, anterior and posterior cingulate cortex, amygdala, and striatum [8–11]. Together, this network supports the perceptual, evaluative, and predictive computations required to interpret others and regulate behavior in complex social worlds [8–13]. The social brain may therefore be shaped by the same relationships it enables animals to navigate.

Consistent with this idea, experimental and observational studies have linked primate social environments to variation in brain structure and function. Macaques housed in larger social groups for more than a year showed greater grey-matter volume in the middle superior temporal sulcus (STS) and rostral prefrontal cortex, and stronger frontal–temporal coupling [14]. More recent work in free-ranging macaques found that the number of direct grooming partners likewise covaried with mid-STS and insular cortex morphology [15]. Human studies similarly link social-network size to brain morphology [16–18] and show that acute isolation elicits social craving with associated midbrain responses [19]. In macaques, social housing can also reversibly alter striatal dopamine D2/D3 receptor availability [20]. Moreover, while previous morphometric studies have emphasized cortical territories, social adaptation is known to require distributed circuitry that extends into the subcortex and brainstem. In macaques, social status covaries with morphology and functional coupling across cortical, sub-cortical, and brainstem systems [21]. Complementary circuit studies in macaques and models of social homeostasis in rodents specifically implicate thalamic, striatal, and brainstem pathways in linking social perception to motivated action and physiological regulation [22–27] What remains unresolved, however, is whether abrupt changes in social contact remodel these distributed systems, whether that remodeling is sustained while the new conditions persist, and whether it reverses when prior conditions return.

Recent work demonstrates that social adaptation depends not only on which brain regions change but also on how they functionally interact as a distributed system. In macaques, observing and participating in social interactions recruits a whole-brain network spanning the STS and medial-lateral prefrontal cortex, operating with limbic structures such as the amygdala [9, 12, 28, 29]. Many of these territories intersect with the default-mode network (DMN)[8], whose anatomy and connectivity are associated with perceived social isolation in humans [30, 31]. A changing social environment may therefore alter not only individual connections but also the embedding of regions within the brain’s global functional architecture. Connectivity manifolds capture this organization by positioning regions according to the similarity of their whole-brain connectivity profiles, allowing shifts in regional embedding to be tracked across conditions [32–38]. Whether an adult social transition reorganizes this functional geometry—and whether those changes recede when prior conditions are restored—remains un-explored.

Here, we followed 13 adult male cynomolgus macaques through three successive phases of single housing, protected-contact paired housing, and restored single housing. Because animals retained visual, auditory, and olfactory contact with familiar conspecifics throughout, pairing introduced sustained proximity to a familiar partner and limited tactile access. At six assessments approximately 3 months apart (two per phase), we acquired structural and resting-state functional MRI and measured hair cortisol as an index of longer-term glucocorticoid exposure [39, 40]. This design allowed us to track the emergence, maintenance, and reversal of neural and physiological responses to the housing transition within the same animals. We found that pairing was associated with grey-matter expansion across several temporal cortical, subcortical, and brainstem territories, functional reorganization of predominantly default-mode and limbic regions, and increased hair cortisol. Within 3 months after single housing resumed, grey-matter and cortisol elevations receded and regional functional changes reversed direction, revealing a reversible component of adult brain architecture that reflects current social conditions.

## RESULTS

All 13 animals completed six longitudinal imaging sessions at approximately 3-month intervals, with two sessions during each of three successive phases: initial single housing (sessions 1-2), protected-contact paired housing (sessions 3-4), and restored single housing (sessions 5-6, Fig. 1A). Voxelwise grey matter, regional functional-manifold measures, and hair cortisol were analyzed using a common longitudinal mixed-effects model with housing condition and centered session as fixed effects and animal as a random intercept (Fig. 1B). In the complete imaging design, housing condition was orthogonal to linear session number, allowing the housing contrast to be estimated separately from a linear trend across visits.

**Fig. 1.**
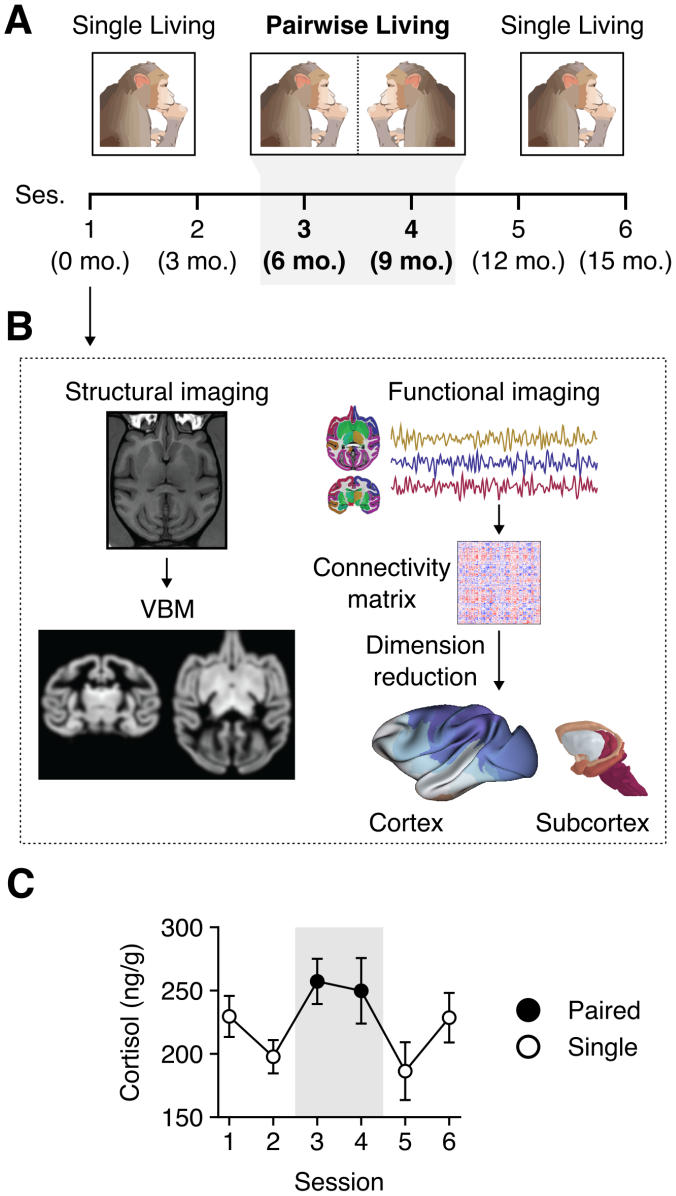
Housing protocol, imaging analyses, and endocrine response. (A) Single–paired–single housing protocol, with two measurement sessions (Ses.) in each phase, separated by ∼3 months. Macaques were initially housed singly (sessions 1-2), then paired in adjacent compartments separated by a wire mesh divider (sessions 3-4), before returning to single housing (sessions 5-6). This design allowed changes associated with the introduction and withdrawal of paired housing to be examined within the same animals. (B) Overview of the structural and functional imaging analyses used across sessions. Left, structural MRI and voxel-based morphometry (VBM) quantified changes in regional grey-matter volume. Right, regional functional MRI time series were used to construct connectivity matrices, followed by dimensionality reduction (diffusion map embedding) to characterize large-scale functional reorganization across cortical and subcortical regions. (C) Hair cortisol concentrations across the six assessments (mean ± SEM). Grey shading and filled symbols indicate paired housing; open symbols indicate single housing. Samples were collected approximately one week before MRI. Individual condition comparisons and exploratory associations are shown in Fig. S8.

The housing transitions were accompanied by changes in hair cortisol, an index of longer-term glucocorticoid exposure [39, 40]. Concentrations were significantly higher during paired housing than during the flanking single-housing phases (*β* = 45.9 ng/g, 95% CI [19.2, 72.6], Wald *z* = 3.37, *p* < 0.001; Fig. 1C and Fig. S8), before declining at the first assessment after single housing resumed. This pattern indicates that the housing transition was accompanied by a sustained shift in the animals’ physiological state, although elevated cortisol does not imply that pairing was aversive [41]. Having established this accompanying endocrine response, we next examined how brain anatomy and functional organization changed across the same housing transitions.

### Paired housing induces reversible grey-matter expansion across distributed brain systems

Using longitudinal voxel-based morphometry, we applied the common longitudinal mixed-effects model at each voxel across the six structural scans from each animal (Fig. 2A). Eleven clusters showed significantly greater grey matter volume during paired housing than during the flanking single-housing phases (voxelwise *p* < 0.001; cluster-corrected *p* < 0.05; table S1). By contrast, no cluster showed significantly lower grey matter volume during paired housing at the same corrected threshold. Six of these clusters were distributed across temporal association cortex in both hemispheres, with additional effects in bilateral posterior cingulate cortex, left striatum, thalamus, and medulla (Fig. 2A; table S1). The concentration of effects in temporal association cortex is consistent with prior links between social group structure and STS morphology [14, 15](Fig. S1), while their extension into posteromedial, subcortical, and brainstem regions indicates a broader anatomical response to the housing transition. Visualizing the data at a more permissive statistical threshold [see 42; uncorrected p < 0.05; Fig. S2], the majority of these effects are embedded within broader clusters that are mirrored bilaterally, with additional clusters observed in regions previously implicated in social processing, including orbitofrontal cortex [9, 10]. This exploratory pattern suggests that the clusters surviving correction formed part of a more broadly distributed structural response.

**Fig. 2.**
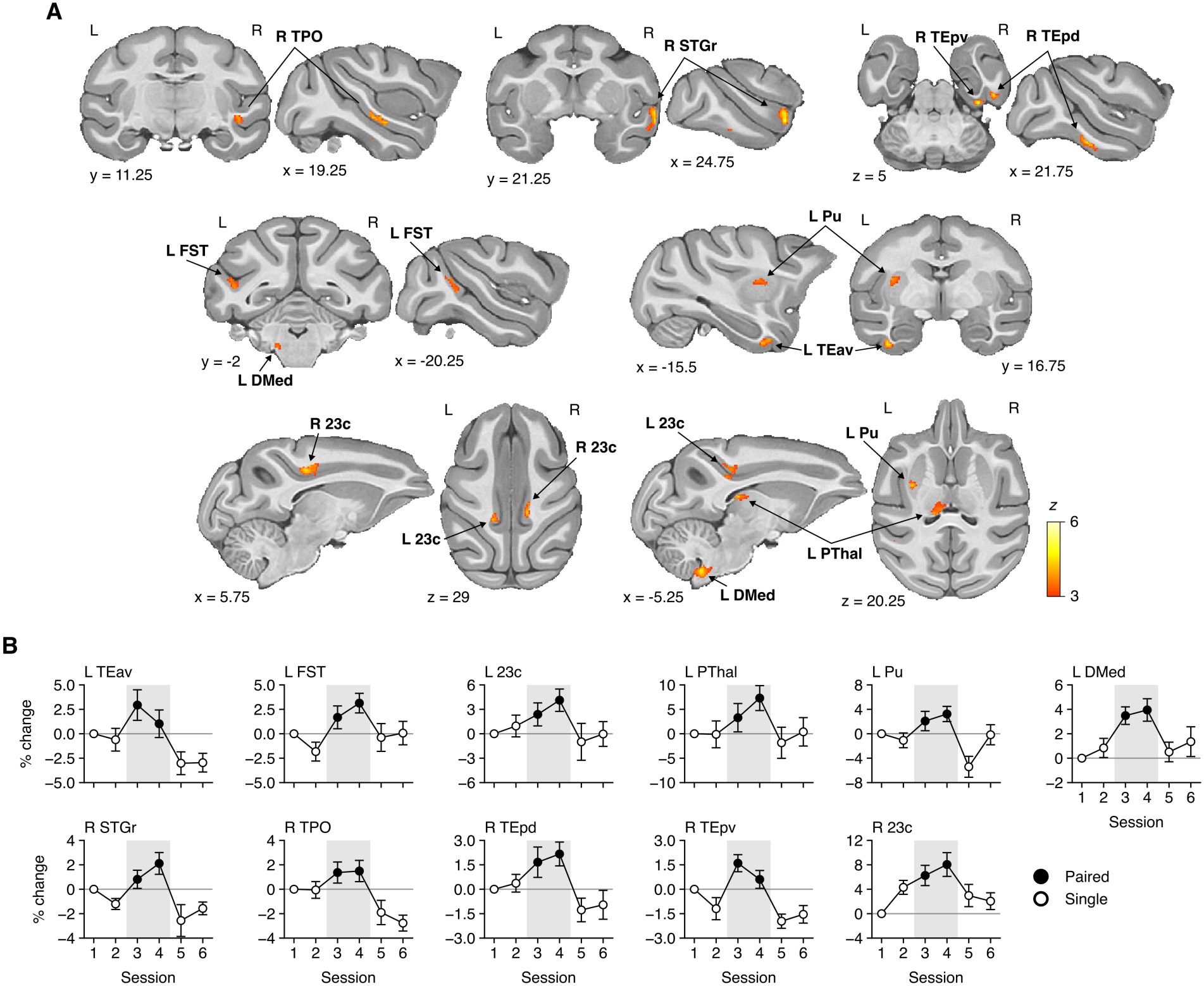
Regional grey-matter expansion during paired housing recedes after single housing resumes. (A) Eleven clusters showing greater grey-matter volume during paired than single housing, identified using voxel-based morphometry (voxelwise *p* < 0.001; cluster-corrected *p* < 0.05). Significant clusters are overlaid on the anatomical template, with colours indicating *z* statistics and slice coordinates given in millimetres in NMT v2 space. Labels identify the corresponding regions; L and R denote the left and right hemispheres. (B) Session-resolved grey-matter changes within each cluster, expressed as percentage change with respect to session 1. Points and error bars show the mean ± SEM. Shading and filled symbols indicate paired housing (timepoints 3–4); open symbols indicate single housing (timepoints 1–2 and 5–6). Across these regions, grey-matter volume increased during paired housing and declined after single housing resumed, consistent with a reversible structural response to the social environment.

To visualize how this expansion unfolded, we plotted mean grey-matter values from each significant cluster across all six imaging sessions (Fig. 2B). Because these clusters were defined by the paired-versus-single housing contrast, the extracted values are presented descriptively and were not used for independent statistical inference. Across clusters, values changed little between the initial single-housing scans (sessions 1-2), increased by the first scan approximately 3 months after pairing began (session 3), and remained elevated at the second paired-housing scan (session 4). Values then declined by the first scan after single housing resumed (session 5) and generally remained at these lower levels at the final scan (session 6), with the extent of return toward initial values varying across regions. Consistent with this pattern, we found that mean changes across the key housing transition intervals were positive from sessions 2 to 3 and negative from sessions 4 to 5 in all 11 clusters (Fig. S3).

To distinguish changes associated specifically with the introduction and withdrawal of paired housing, we divided single housing into Pre (sessions 1-2) and Post (sessions 5-6) and computed pairwise contrasts among the Pre, Paired, and Post phases, using the same voxel-wise and cluster-corrected thresholds (Fig. S4). Of the 11 original clusters, three spatially overlapped only with the Pre-versus-Paired contrast map, five only with the Post-versus-Paired map, and three with both maps. None overlapped with significant Pre-versus-Post differences, consistent with a return toward pre-pairing grey-matter volumes, particularly in regions showing both emergence and reversal. Together, the trajectories and phase comparisons support reversal of the pairing-associated expansion, with the extent of return toward initial grey-matter values varying across regions.

### Mapping housing-related changes in functional organization

We next examined how distributed functional organization changed across the housing transitions. For each animal and resting-state fMRI session, we extracted mean BOLD time series from a whole-brain macaque parcellation spanning cortical, subcortical, and brainstem structures (see Methods). We represented the functional relationships among these 316 regions (264 cortical and 52 subcortical) in a 316 × 316 covariance matrix.

Stable individual signatures can dominate functional connectivity matrices and obscure smaller state-dependent changes [34, 43, 44]. In our data, animal identity accounted for 19.9% of variation, and scans grouped predominantly by animal rather than housing condition (Fig. 3A). We therefore applied Riemannian centering to re-express each animal’s six covariance matrices around a common group mean while retaining within-animal variation across sessions [45]. After centering, nearest-neighbor identification accuracy fell from 66.7% to 0% (chance: 6.5%), whereas the proportion of variation associated with housing condition remained similar (1.2% before centering and 1.5% afterward). These centering diagnostics indicate reduced stable individual signatures while retaining a similar housing-related component, consistent with previous applications of this approach [34, 36, 37, 46].

**Fig. 3.**
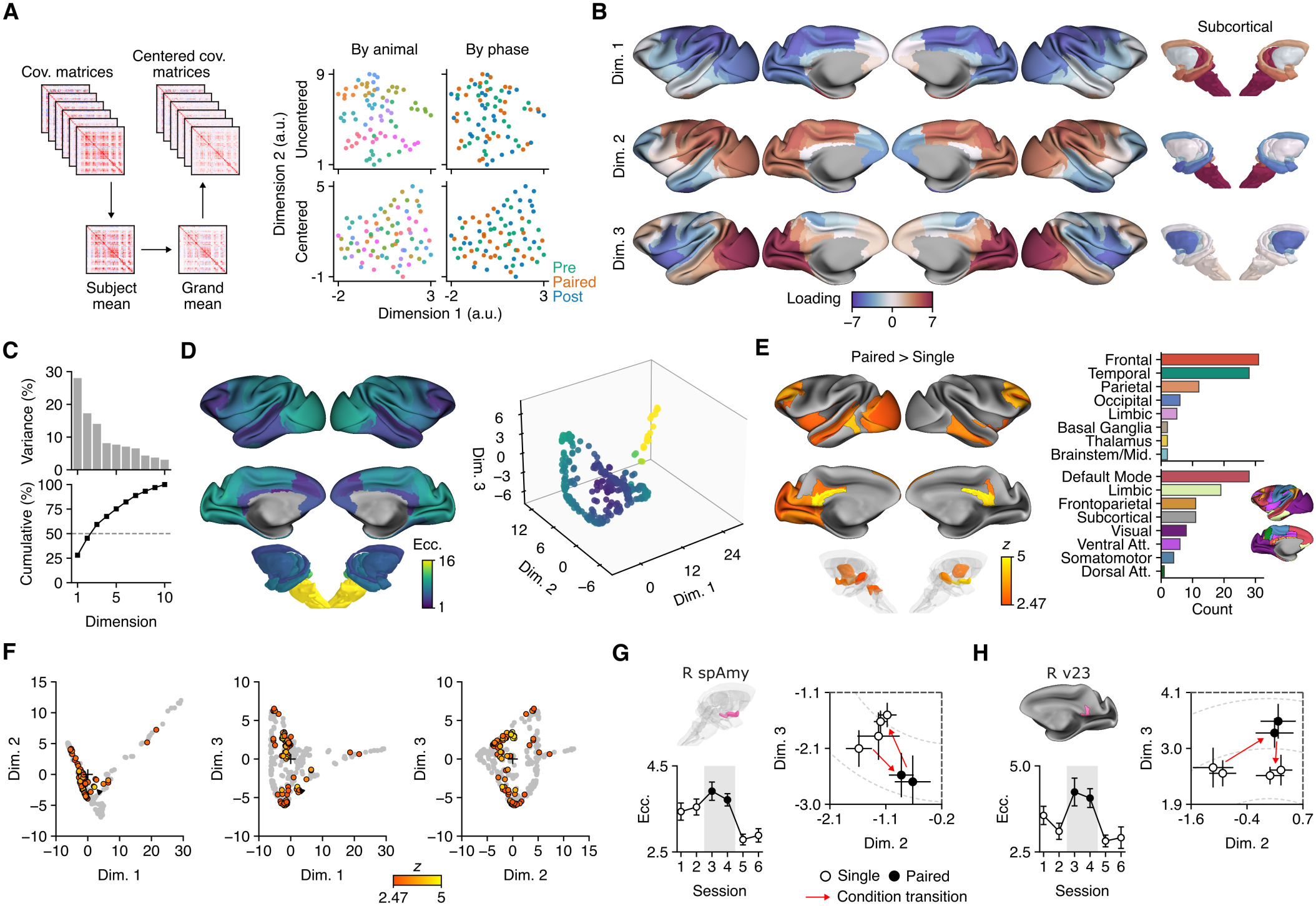
Paired housing increases regional differentiation within the functional-connectivity manifold. (A) Riemannian centering aligns animal-specific mean covariance matrices to a common grand mean. UMAP visualizations of all 78 matrices (13 animals × 6 sessions; one point per scan) both before (top) and after (bottom) centering, coloured by animal (left) or housing phase (right). UMAP is shown for visualization purposes only. (B) The first three dimensions of the template connectivity manifold, computed using the grand mean across all matrices, displayed on cortical and subcortical surfaces. colours indicate regional dimension loadings. (C) Individual and cumulative contributions of the first ten diffusion-map dimensions normalized by the sum of the ten computed eigenvalues; the first three account for 59.3%. (D) Template manifold eccentricity displayed anatomically and in three-dimensional manifold space. Each point represents a region; colours indicate distance from the embedding origin. (E) Regions with greater eccentricity during paired housing than during the flanking single-housing phases in the longitudinal model, with linear session and housing condition as fixed effects and animal as a random intercept. Wald *z* shown for the 88 of 316 regions that survived FDR correction (*q* < 0.05); all significant effects were positive. Counts of significant regions by anatomical division (top right) and functional-network assignment (bottom right). Counts summarize the anatomical and network distribution of significant effects. (F) Location of the significant regions in three-dimensional manifold space. Significant regions are highlighted with the colour scale in (E), and non-significant regions are shown in grey. (G, H) Eccentricity changes for two exemplar regions (right subpallial amygdala [spAmy]; right area v23); regions are shown in pink in the subcortical and cortical renderings. Line plots show mean ± SEM eccentricity across sessions (bottom). Zoomed-in point plots show the region’s mean ± SEM manifold embedding projected onto the second and third dimensions (right); red arrows indicate transitions into and out of paired housing (filled circles). grey dashed arc lines indicate distance from manifold origin (1-unit spacing).

We next used multivariate distance matrix regression (MDMR) [47–49] to test whether housing condition was reflected in whole-brain connectivity patterns. This approach tests whether differences among whole-connectome patterns are explained by the experimental design, without testing individual connections separately. We modeled correlation distances (1 − *r*) among centered connectivity matrices as a function of housing condition, accounting for animal identity and the linear effect of session. Significance was assessed by permuting condition labels within each animal (see Methods). We found that housing condition explained a significant component of whole-connectome variation (pseudo-*F* = 2.07, partial *R*^2^ = 0.033, permutation *p* = 0.008). We next examined where functional organization changed and how these regional effects unfolded across the housing transitions.

### Default-mode and limbic regions reversibly shift within the functional-connectivity manifold during paired housing

To examine regional changes in functional organization, we used manifold learning to embed regions in a low-dimensional connectivity gradient space, where regions with similar whole-brain connectivity profiles lie near one another [32, 34, 36, 38, 50]. The first three dimensions of the template manifold accounted for 59.3% of the sum of the ten computed diffusion-map eigenvalues (Fig. 3B,C). Dimension 1 primarily distinguished subcortical and brainstem regions from cortex, with positive loadings in the former and negative loadings across much of visual and somatomotor cortex. Parts of association cortex occupied intermediate positions, with near-zero or weakly positive loadings. Dimension 2 more clearly separated visual and somatomotor systems from the default-mode network (DMN) and other association territories, closely resembling the sensory-to-transmodal organization described in previous work [32]. Along this axis, brainstem regions lay toward the sensory–motor end and hippocampal and amygdalar regions toward the DMN end. Dimension 3 further distinguished the sensory–motor systems, separating visual cortex from lateral somatomotor regions.

To quantify housing-related shifts within this functional architecture, we calculated each region’s manifold eccentricity, defined as its distance from the embedding origin across the retained dimensions [33–38] Greater eccentricity indicates a more differentiated whole-brain connectivity profile within this space (Fig. 3D). Housing-condition effects on regional eccentricity were tested using the longitudinal mixed-effects model described above. To relate these geometric shifts to changes in coupling, we subsequently examined within- and between-system connectivity.

Paired housing increased manifold eccentricity in 88 of 316 regions, with no significant decreases (FDR *q* < 0.05; Fig. 3E; table S2). More than half of the responsive regions belonged to the DMN and limbic network (28 and 19 regions, respectively; 47 of 88, 53%), spanning posteromedial, prefrontal, and temporal cortex. Subcortical and brainstem effects included the bilateral amygdala and thalamus, left hypothalamus, dorsal and lateral pons, and globus pallidus. Temporal cortex accounted for 28 of the 88 responsive regions (32%), including regions along the superior and inferior temporal cortex. The prominence of the temporal association cortex echoed the structural findings, with functional effects encompassing territories within macaque systems for processing faces, bodies, social interactions, and conspecific vocal signals [12, 28, 51, 52].

To illustrate the time course of these effects, we examined session-by-session estimates in the right subpallial amygdala and right area v23 as illustrative subcortical and cortical regions, respectively. Both showed elevated eccentricity during paired housing, followed by a decline after single housing resumed (Fig. 3G,H). Note that these regions’ trajectories are shown for descriptive purposes rather than a time-resolved analysis of changes in eccentricity.

The eccentricity effects were largely retained after accounting for regional BOLD variance and across alternative analytical choices. Regional BOLD variance showed no significant housing-condition effect in any of the 316 regions (Fig. S5A), and 80 of the 88 eccentricity effects remained significant when variance was included as a covariate (Fig. S5B,C). A repeated-measures ANOVA recovered 58 of the 88 significant regions, with regional condition statistics nearly perfectly correlated with those from the mixed-effects model (*r* = 0.999; Fig. S6A). The spatial pattern also remained stable when including a larger number of diffusion-map dimensions, different row-retention thresholds, and alignment templates (Fig. S6B–D). Condition effects were strongly correlated between left–right homologues (*r* = 0.79; Fig. S7B). Together, these analyses identify a housing-related re-organization of functional architecture that was consistent across analytical choices. Regional eccentricity increased during pairing, remained elevated across the paired-housing sessions, and receded after single housing resumed, paralleling the emergence, maintenance, and recession of the structural response.

### Complementary connectivity changes underlie manifold differentiation

Manifold eccentricity identifies regions with altered whole-brain connectivity profiles but does not reveal which connections account for those shifts. The right subpallial amygdala and right area v23 illustrate this distinction: both increased in eccentricity during pairing, yet their connectivity changes followed very different patterns (Fig. 4A). Accordingly, for each of the 88 responsive regions, we profiled whole-brain connectivity change directly by taking the difference in each connection’s Fisher *z* correlation between the paired and single housing conditions. Clustering regions by the similarity of these whole-brain profiles revealed two main modes of reorganization, with both the silhouette and Calinski– Harabasz criteria favoring a two-cluster solution (Fig. 4B).

**Fig. 4.**
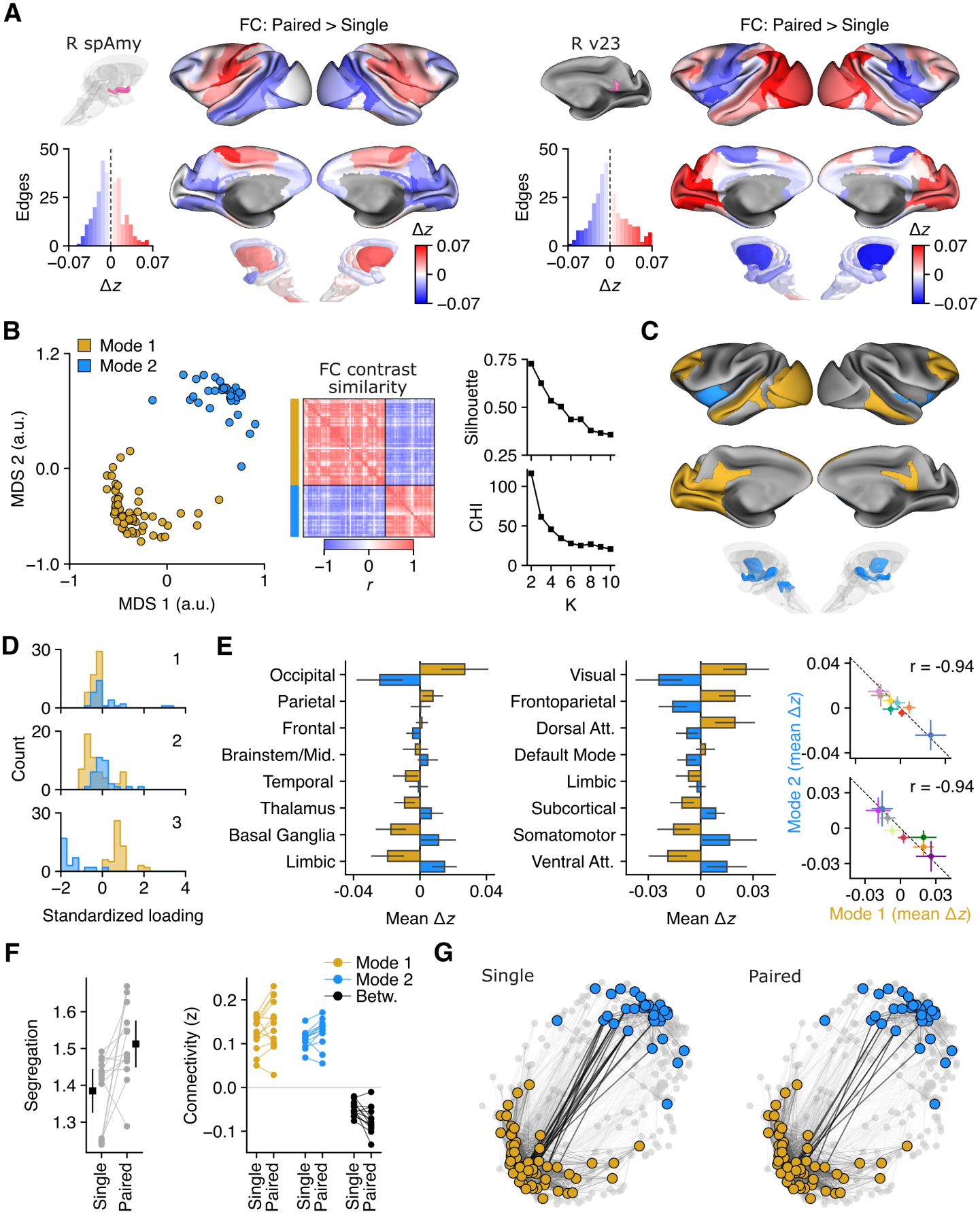
Two complementary connectivity-change modes accompany housing-related manifold differentiation. (A) Whole-brain connectivity change distributions and maps for right subpallial amygdala and right area v23 (denoted in pink). Histograms and map colours indicate edgewise differences in functional connectivity (Fisher *z*) between paired and single housing, with positive values denoting greater connectivity during pairing. (B) Organization of the 88 responsive regions by the similarity of their whole-brain connectivity-change profiles. Multidimensional scaling displays profile dissimilarities, with one point per region; the matrix shows correlations between profiles ordered by mode. Silhouette and Calinski–Harabasz cluster criteria are shown for k = 2–10. Yellow and blue indicate Modes 1 and 2, respectively. (C) Anatomical distribution of the cortical-associative and limbic-subcortical modes. (D) Distributions of regional template manifold loadings along the first three gradients, separated by mode. (E) Mean connectivity change (Fisher *z*) for each mode, summarized by target anatomical division (left) and functional-network category (middle). Error bars indicate the standard deviation across regions contributing to each mode’s category. Scatterplots (right) compare the two modes’ category-level means, with correlations of *r =* − 0.94 and *r =*− 0.94, respectively. (F) Mode segregation (left) and within-mode and between-mode connectivity (right) during single and paired housing, computed from per-session Fisher *z* connectivity. Segregation is the system-segregation index (see Methods). Each animal’s single- and paired-housing values are the means across its single-housing sessions (sessions 1-2 and 5-6) and paired-housing sessions (sessions 3-4), respectively. Points represent animals, and lines connect each animal’s condition means. Left: mean ± 95% CI (black). Right: coloured by connectivity type (within Mode 1, within Mode 2, and between modes). Because mode membership was derived from housing-related connectivity changes in the same data, these summaries describe the identified response rather than provide an independent test of segregation. (G) Connectivity diagrams for single and paired housing. Responsive regions are fixed at the MDS coordinates and are coloured by mode assignment. Black edges indicate between-mode connectivity; all other brain areas and connectivity are shown in faded grey (top 20% of connections shown). Between-mode connectivity falls under pairing, reflecting increased segregation.

The cortical-associative mode (Mode 1, yellow; 55 regions) spanned temporal, prefrontal, posteromedial and visual cortex. The limbic-subcortical mode (Mode 2, blue; 33 regions) encompassed insular and limbic territories together with subcortical structures, including the amygdala and hypothalamus (Fig. 4C). Anatomically, this broad division resembles prior distinctions in macaques between temporal–prefrontal circuits implicated in social-information processing and limbic-subcortical circuits associated with social status and state regulation [10, 12, 14, 21]. The two modes also occupied distinct portions of the whole-brain manifold, particularly along its second and third dimensions (Fig. 4D). The two modes expressed broadly opposing patterns of connectivity change (Fig. 4E). During paired housing, cortical-associative regions (Mode 1) showed stronger coupling with frontoparietal, visual, and dorsalattention networks, but reduced coupling with somatomotor, ventral attention, limbic and subcortical regions. Limbic-subcortical regions (Mode 2) generally showed the converse pattern for these same targets. Across the functional-network and subcortical categories, the modes’ mean connectivity-change profiles were strongly negatively correlated (*r* = − 0.94), with a similar opposition across anatomical divisions (*r* = − 0.94).

We next used within- and between-mode connectivity and system segregation to describe the response of these data-derived modes (Fig. 4F). During paired housing, connectivity increased within both modes and became more negative between them, yielding greater functional segregation (Fig. 4F,G). Together with the manifold findings, these observations describe stronger internal coupling and greater separation between cortical-associative and limbic-subcortical systems during pairing.

## DISCUSSION

By following the same adult macaques through the introduction and withdrawal of paired housing, we identified reversible changes in brain anatomy and the organization of distributed functional systems. Grey-matter expansion spanned cortical, subcortical, and brainstem territories, with most effects concentrated in temporal association cortex. Functional reorganization pre-dominantly involved default-mode and limbic regions, and was accompanied by greater segregation between cortical-associative and limbic-subcortical systems. Hair cortisol also increased during paired housing, showing that the neural changes were accompanied by a measurable shift in longer-term endocrine state. The responses emerged within months, were maintained over the duration of paired housing, and receded by the first imaging session approximately 3 months after single housing resumed. These findings reveal a form of biological embedding in which changes to mature brain architecture are sustained while social conditions persist, yet recede when original conditions are restored.

### Changing social conditions reshape primate brain architecture across scales

Temporal association cortex was a prominent site of both grey-matter changes and functional reorganization across the housing transitions. This convergence builds on findings from laboratory and free-ranging macaques linking temporal cortex morphology to social group size and the number of direct grooming partners [14, 15]. These territories encompass systems for perceiving faces and bodies, interpreting social interactions, and processing conspecific vocalizations [9–12, 51–55]. Our longitudinal findings show that this anatomical response can emerge within months, persist during paired housing, and recede after single housing resumes, revealing its sensitivity to the continuation and abrupt withdrawal of close social contact.

The functional findings extend this anatomical response to the relationships among distributed brain systems. Our whole-connectome analysis identified housing-related changes in connectivity organization and our manifold analyses localized this reorganization largely to DMN and limbic territories, including bilateral posterior cingulate cortex. These posteromedial regions form a central component of the primate DMN [8, 56], whose organization in humans covaries with the perceived social environment, including loneliness [8, 30, 31]. During pairing, this posteromedial system participated in a cortical-associative mode that became more segregated from a complementary limbic-subcortical mode. Coupling increased within both modes and decreased between them, indicating that stronger internal connectivity accompanied greater differentiation between systems. Previously reported increases in temporal-frontal coupling with social-group size [14] may therefore reflect one aspect of the broader reorganization of association-cortical connectivity observed here during paired housing. Regional eccentricity subsequently declined after single housing resumed, linking these changes in functional organization to the broader reversible response.

Grey-matter expansion in the putamen, mediodorsal thalamus, and dorsal medulla, together with functional reorganization in the amygdala and hypothalamus, extends the response to systems linking social information with action and physiological regulation. Previous macaque morphometry associated social rank with variation in striatal and brainstem anatomy [21]. Here, repeated imaging shows that these territories also change within individuals across the introduction and withdrawal of paired housing. Striatal circuits contribute to social action and reward processing [23], while the amygdala evaluates socially relevant information and helps guide responses to other individuals [9, 57, 58]. Recent rodent studies further illustrate how plasticity in thalamocortical circuits can modify social behavior and status [24, 27]. The hypothalamus and medullary systems, in turn, help translate information about bodily state and environmental demands into autonomic and neuroendocrine responses [59, 60]. Together with the accompanying cortisol changes we observed, this distribution suggests that adaptation to changing social contact extends across systems for social evaluation, motivated action, and physiological regulation, consistent with recent models of social homeostasis [26].

### Reversible remodeling in the mature primate brain

Studies of early social deprivation in humans and macaques have emphasized the persistent neural and behavioral consequences of disrupted developmental experience [61–64] That the changes observed here emerged and receded in fully mature animals is therefore central to their interpretation. Adult human neuroimaging has documented training-related structural changes that partially regress after training ends [65], as well as hormone-associated variation in grey-matter volume and cortical thickness across the menstrual cycle [66, 67]. Our findings extend this literature by showing that structural changes associated with a sustained alteration in social housing can also recede when prior conditions return, alongside shifts in distributed functional organization. This continuing responsiveness may be particularly consequential across primates, including humans, whose relationships and social-network positions change throughout adulthood and can be abruptly reorganized by major environmental disruptions (e.g., natural disasters, pandemics)[6, 68–71]

### Distributed remodeling follows increased proximity and tactile

The form of social contact the animals experienced is central to interpreting the neural remodeling we observed. Throughout the study, animals remained in a shared room with continuous visual, auditory, and olfactory access to familiar conspecifics. Pairing changed the immediacy of that contact by bringing a familiar partner into sustained close proximity and allowing limited tactile interaction through a wire-mesh divider. This protected-contact arrangement permitted closer engagement while reducing the risk of aggression and injury associated with unrestricted contact between adult males, consistent with established macaque-management practices [72–74]. Animals could therefore interact at close range but could neither groom one another nor make unrestricted physical contact. Despite these constraints, we found that closer access to an already familiar partner was accompanied by remodeling that extended from local grey matter changes to the reorganization of distributed functional systems.

Closer engagement, however, need not be uniformly affiliative. Macaques can express affiliative, agonistic, and tension-related behaviors across barriers, and the physiological consequences of those social interactions can vary with relationship quality [72–74]. The studies differ in species, sex composition, and contact arrangements, which limits direct extrapolation to our male cynomolgus cohort. The increase in hair cortisol that we observed indicates greater glucocorticoid exposure during pairing but does not establish whether the experience was rewarding, demanding, adverse, or some combination of these [41, 75]. The neural changes may therefore reflect adaptation to the demands of sustained proximity—including carefully monitoring a partner, predicting their actions, and regulating one’s own responses—alongside opportunities for affiliation. These findings suggest that adult primate brain architecture adjusts to the informational and regulatory demands of closer social engagement [9, 11], even when physical interaction remains limited.

### Methodological considerations

The fixed A–B–A housing sequence is both a central strength and the principal constraint of our design. Six measurements spanning the introduction, maintenance, and withdrawal of paired housing allowed each animal to serve as its own longitudinal reference, but the absence of a contemporaneous control prevents full separation of housing effects from other influences associated with study period. Because the paired-housing scans fell midway between the initial and final single-housing scans, a change progressing at a constant (linear) rate across scans would not produce the observed paired-versus-single difference. More complex changes (non-linear) over time cannot be excluded, but an alternative account would need to explain the relative stability between measurements within phases, the opposing changes following the introduction and withdrawal of paired housing, and the return toward initial anatomical, functional, and endocrine states. Together, these trajectories argue against a simple effect of elapsed time and support a state-dependent response to the housing transition.

The scope of the housing transition should also be defined precisely. Protected-contact paired housing increased proximity to a familiar partner, introduced limited tactile access, and changed enclosure size and configuration. Because these features changed together, their individual contributions cannot be separated in the present design. Nevertheless, our STS expansion closely matches the grey-matter effect reported by Sallet et al., whose larger social groups did not have larger enclosures [14], and overlaps the foci identified by Testard et al. in free-ranging macaques [15](Supplementary Fig. 1). This anatomical convergence across distinct social and physical environments strengthens the evidence for socially driven structural plasticity in the STS. However, because interactions across the mesh divider were not quantified systematically, we cannot determine how the neural changes relate to specific social behaviors, relationship quality, or affective valence. Future studies combining repeated imaging with detailed behavioral measurements and experimentally varied forms of contact will be useful in determining which specific aspects of social access drive the observed remodeling.

Further questions concern how broadly these findings generalize, how quickly the changes unfold, and which biological processes they reflect. First, the cohort included only males. Sex differences in wild cynomolgus social organization, including male dispersal and female residence in natal groups [76, 77], motivate testing whether the magnitude or distribution of neural remodeling, or the endocrine response to protected contact, differs in females. The five dyads and one triad offered limited scope to test effects of partner number or social-network complexity, and responses among animals sharing a social unit may be correlated. Second, scans approximately three months apart established that remodeling was clearly detectable within months of each housing transition, but could not resolve when changes began, how they progressed, or how rapidly they receded. Third, T1-weighted morphometry reflects multiple tissue properties and cannot identify the cellular processes underlying grey-matter expansion. Resolving these processes will require longitudinal microstructural imaging and ultimately histological validation [78, 79]. Finally, the structural, functional, and endocrine responses were analyzed separately. Their parallel trajectories link each to the same housing transition but do not establish whether one response influenced another or whether they shared an underlying mechanism.

### A reversible imprint of the social environment

Within months, we found that changing housing conditions were accompanied by remodeling of local grey matter, reorganization of distributed functional systems, and altered endocrine state, with several of these responses receding after single housing resumed. These findings reveal a reversible component of the biological imprint of social experience on the mature primate brain, suggesting that adult brain architecture does not simply retain a fixed record of past conditions but remains responsive as those conditions change. Given the shared features of human and macaque brain organization, these findings offer a window onto how human brain architecture might respond to major social transitions associated with aging, illness, institutional confinement, or societal disruption. The question, then, is not only how the adult brain changes when social conditions are altered, but which of those changes persist when familiar conditions return.

## MATERIALS AND METHODS

### Subjects

Thirteen adult male Mauritian-origin cynomolgus macaques (*Macaca fascicularis*) participated in the study. At study onset, animals were 6–11 years old (mean = 7.5 years) and weighed 6.3–12.8 kg. The cohort comprised animals available for the complete longitudinal protocol; no corresponding female cohort was available at the time. All animals were unrelated and had no prior exposure to the social-housing manipulation. They were housed at the Centre for Neuroscience Studies, Queen’s University (Kingston, Ontario, Canada), under the care of a laboratory animal technician and the institutional veterinarian. All experimental and animal-care procedures were approved by the Queen’s University Animal Care Committee and conducted in accordance with Canadian Council on Animal Care guidelines.

### Experimental design

Animals underwent a longitudinal A–B–A housing protocol comprising an initial single-housing baseline (Phase 1), approximately six months of protected-contact paired housing (Phase 2), and approximately six months of restored single housing (Phase 3). Before the first imaging session, animals had been singly housed for approximately nine months, providing a stable baseline period. Structural and resting-state functional MRI were acquired at sessions 1–6 (two per phase), with successive sessions separated by an interval of 106 ± 12 days (mean ± min/max across the five intersession intervals; Fig. 1A). The two initial sessions therefore provided a pre-pairing reference under established single-housing conditions. Within each scanning window, one to three animals were scanned per day, with animal order preserved across sessions; each scanning window spanned 9–17 days. The fixed A–B–A sequence was selected to test whether neural and endocrine responses emerged during paired housing and receded after single housing resumed.

Partner assignment was based on veterinary and behavioral assessments of compatibility conducted by experienced animal-care staff in consultation with the institute veterinarian. The five dyads and one triad were composed primarily according to individual temperament, with the aim of forming compatible social units while minimizing the risk of aggression or injury during the protected-contact housing phase (see also *Dominance assessment*, below).

### Housing manipulation

Throughout all three housing phases, animals remained in a large shared room with continuous visual, auditory, and olfactory access to familiar conspecifics. The manipulation therefore varied proximity to a familiar partner and the opportunity for limited tactile contact, rather than the presence or absence of social stimulation. During the initial and final single-housing phases, animals occupied individual, non-contiguous stainless-steel cages measuring 2.6 m × 1.3 m × 1.3 m, positioned at least 1 m apart. This separation prevented physical contact between animals. During the initial phase, future protected-contact partners occupied neighboring cages.

During paired housing, animals occupied adjoining pens measuring 2.0 m × 1.5 m × 2.5 m per animal. Adjacent pens were separated by a wire-mesh panel measuring 53-cm × 182.5-cm, with 3-cm × 3-cm apertures. This arrangement provided continuous opportunities for close-range interaction and permitted limited tactile contact through the mesh, while preventing unrestricted physical contact and social grooming.

Animals were organized into five dyads and one triad. In the triad, the animal occupying the central pen had direct protected contact with both partners, whereas the animals in the outer pens had direct protected contact only with the central animal. Social-unit membership remained fixed throughout the paired phase, including during routine cleaning.

### Dominance assessment

Animal-care staff considered dominance-related behavior when forming compatible social units, generally pairing males perceived as more dominant with less dominant partners and avoiding, where possible, pairing two strongly dominant individuals. To assess relative dominance and its stability across the housing transitions, experienced observers rated each animal on the seven-point Dominant item of the macaque personality questionnaire [80]. Three observers provided ratings during each of the first two housing phases, and two new observers provided ratings during the final phase. Ratings were averaged within animal and phase.

Mean ratings placed the same animal higher across all three phases in four of the five dyads. In the fifth dyad, the same animal was rated higher during the first two phases, with equal mean ratings during the final phase. Ordering within the triad varied. Individual raters generally supported the consistent ordering within dyads: of 40 comparisons across dyads and phases, 27 supported this ordering and nine were ties. Two comparisons opposed the ordering; the remaining two came from one dyad during the final phase, in which the observers favored different animals.

Across the full cohort, ratings were concordant across phases (Kendall’s W = 0.66, *χ*^2^(12) = 23.65, *p* = 0.023; pairwise Spearman *ρ* = 0.47–0.57). Baseline ratings also correlated with those provided by the new observers during the final phase (*ρ* = 0.57). Together, these assessments support broadly stable relative dominance within the dyads across the housing transitions, with greater variability in the triad.

### Analysis overview and statistical framework

The structural and functional analyses tested complementary aspects of the housing response. Voxel-based morphometry tested whether local grey-matter volume tracked housing condition. Functional analyses began with multivariate distance matrix regression, an omnibus test of whether housing condition was expressed in whole-brain connectivity, before mani-fold analysis localized that reorganization to individual regions. Connectivity-change profiles and mode summaries then described the organization of the detected functional response. Hair cortisol provided a separately measured endocrine index.

Except where otherwise specified, structural, regional functional, and endocrine outcomes were evaluated using a common mixed-effects model. For a measurement *y* from animal *i* at session *t*:

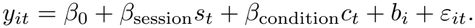

Here, *s*_*t*_ indexes centered session number, *c*_*t*_ codes protected-contact paired housing, and *b*_*i*_ is a random intercept for animal.

In the complete imaging design, the housing vector [0, 0, 1, 1, 0, 0] is orthogonal to linear session number. The condition term therefore estimates the paired-housing difference after accounting for linear trend effects. Three hair-cortisol observations were missing because of insufficient hair regrowth (see below), so exact orthogonality does not hold for that outcome; the mixed model included both session and condition and accommodated the incomplete data.

### MRI preparation and acquisition

The neuroimaging data were acquired using a 3T MRI scanner (MAGNETOM Prisma fit, Siemens, Erlangen, Germany) and a custom 24-channel macaque head coil (Robarts Research Institute, University of Western Ontario). Anesthesia was induced with ketamine (6 mg/kg, intramuscular) and dexmedetomidine (DEX; 4.5 ^*µ*^g/kg, intramuscular), with glycopyrrolate (0.013 mg/kg, intramuscular). Animals were intubated, and a catheter was placed in the saphenous vein. During resting-state acquisition, anesthesia was maintained with intravenous DEX (4.5 ^*µ*^g/kg/hr) and isoflurane at vaporizer settings of 0.6–0.7%, delivered with 100% oxygen at 1.5 l/min. The maintenance regimen was based on established macaque functional-imaging protocols using continuous DEX with low-dose isoflurane [81, 82]. Animals breathed spontaneously. Peripheral oxygen saturation, heart rate, electrocardiogram, and respiration were monitored throughout each session, with intermittent non-invasive blood-pressure measurements. Monitoring supported animal care during acquisition; digitized physiological waveforms were not archived for subsequent analysis, and end-tidal CO_2_ was not recorded. Body temperature was maintained using an MRI-safe ConRad thermal blanket (DRE Veterinary). Anesthetic preparation and acquisition order were held as constant as practicable across sessions.

T1w images were acquired using a 3D Magnetization Prepared Rapid Acquisition Gradient Echo (MPRAGE) sequence (0.5 mm isotropic, FOV = 128 × 128 × 112 mm, matrix = 256 × 256 slices per slab = 224, transversal orientation, readout direction of inferior (I) to superior (S), phase oversampling = 15%, averages = 3, TR = 2200 ms, TE = 2.23 ms, TI = 900 ms, flip-angle = 8°, bandwidth = 270 Hz/pixel, echo spacing = 6.5 ms, no fat suppression, GRAPPA = 2, turbo factor = 224 and pre-scan normalization). The total acquisition time for the T1w scan was 17 min 31 s. Within each session, images were acquired in the order: localizer, T1w, the first resting-state run and its field maps, the second restingstate run and its field maps.

Pairs of spin-echo echo-planar images with opposing phase-encoding directions (right-to-left and left-to-right) were acquired immediately after each resting-state run to estimate susceptibility-induced distortion (1.5 mm isotropic, FOV = 148 mm, 52 slices, matrix = 98 × 98, TR = 8000 ms, TE = 66 ms, flip-angle = 90°, bandwidth = 1760 Hz/pixel, echo spacing = 0.71 ms, EPI factor = 98, phase partial Fourier = 7/8, no parallel imaging, fat saturation, 3 measurements per phase-encoding direction). Each field-map acquisition lasted 32 s.

Resting-state functional images were acquired using a multiband gradient-echo echo-planar imaging sequence (1.5 mm isotropic, FOV = 148 mm, 52 slices, matrix = 98 × 98, TR = 1150 ms, TE = 33.8 ms, flip-angle = 45°, multiband acceleration factor = 4, no in-plane parallel imaging, bandwidth = 1760 Hz/pixel, echo spacing = 0.71 ms, EPI factor = 98, phase partial Fourier = 7/8, fat saturation). Two runs of 2,350 volumes were acquired in each session, one with right-to-left and one with left-to-right phase encoding (45 min 11 s per run; approximately 90 min of resting-state data per animal per session), each followed immediately by its pair of spin-echo field maps. The two runs were concatenated after pre-processing to yield a single time series per session. Animals were anesthetized throughout and no stimulus was presented during acquisition.

### Structural MRI preprocessing and quality control

Longitudinal voxel-based morphometry [83] was performed using Advanced Normalization Tools (ANTs v2.4.3; [84]), based on established ANTs longitudinal procedures [85, 86]. For each animal, the six session T1w images were brain-extracted using pre-computed brain masks with manual corrections, rigidly aligned, and used to construct an unbiased within-subject longitudinal template via antsMultivariateTemplateConstruction2. Each within-subject template was then nonlinearly registered to the NIMH Macaque Template version 2.0 (NMT v2; [87, 88]). This hierarchical approach reduced dependence on any individual session and established correspondence both within animals and across the group.

The within-subject templates were segmented into four classes (cortical grey matter, subcortical grey matter, white matter, CSF) using Atropos [89], initialized with tissue priors derived from the binary tissue masks supplied with NMT v2 (smoothed with a 1mm FWHM Gaussian kernel and aligned to each within-subject template). Each session’s T1w image was then denoised, N4 bias-field corrected [90], and nonlinearly registered to its within-subject template (ANTs SyN)[91]. The template tissue posteriors were propagated to session space to initialize per-session segmentation. Local grey-matter volume was quantified by modulating the session’s cortical and subcortical grey matter segmentations using the Jacobian determinant of the session-to-template warp. Combined modulated grey-matter maps were then normalized to NMT v2 space, and underwent Gaussian smoothing (1mm FWHM).

All source images, brain masks, tissue segmentations, within-subject templates and spatial-normalization results were inspected visually for image quality and processing accuracy. No scans were excluded from analysis; registration accuracy was high for all sessions and animals (normalized cross-correlation [NCC] = 0.88 ± 0.02 [mean ± SD]; n = 78). To account for residual variation in registration quality, session NCC was included as a co-variate in voxel-based morphometry (VBM) models.

### Primary voxel-based morphometry analysis

Whole-brain voxelwise longitudinal models were implemented in Julia using MixedModels v4.25, which tested whether grey-matter volume tracked the housing manipulation using the common longitudinal model defined above. Local grey-matter volume was used as the outcome and the condition term coding the paired-housing sessions; the primary contrast compared protected-contact paired housing with single housing (two-sided test). Statistical maps were thresholded at a voxelwise level of two-tailed *p* < 0.001 and cluster-corrected to *p* < 0.05 within a whole-brain NMT v2 mask. Cluster correction used AFNI’s 3dClustSim (version 25.2.18; [92, 93]), with spatial autocorrelation estimated from model residuals (via 3dFWHMx), yielding a minimum cluster extent of 140 voxels (17.5mm^3^).

Clusters were anatomically labeled by the atlas region at each cluster’s peak voxel, using the Cortical Hierarchy Atlas of the Rhesus Macaque (CHARM; [87]) and the Subcortical Atlas of the Rhesus Macaque (SARM; [94]), supplemented by visual inspection in NMT v2 space. Co-ordinates, volumes, peak statistics, and anatomical labels for all significant clusters are reported in table S1. To visualize the temporal profile of the detected group effect, we extracted a session-resolved estimate for each significant cluster as the mean modulated grey-matter volume within it, computed separately for each animal and session.

### Resting-state fMRI preprocessing and quality control

Resting-state data were preprocessed using FSL v6.0 [95], and ANTs v2.4.3 [84]. Gradient nonlinearity distortions in each BOLD run and its single-band reference was corrected using the Human Connectome Project gradunwarp tool [96, 97]. The initial 10 volumes of each run were discarded to allow signal stabilization, yielding 2 × 2,340 = 4,680 volumes per session in 71 of the 78 scans (four scans contain 5,380 volumes, and three scans contain 3,990–4,343 volumes). Head motion was corrected by rigid-body registration of the BOLD time series to the single-band reference using MCFLIRT [98]. Susceptibility distortions were corrected using FSL topup with opposing phase-encoded (LR/RL) acquisitions [98, 99], followed by synthetic-image-based refinement. Synthetic BOLD targets were generated from T1-weighted images using radial-basis-function regression with interaction terms, spatial blurring, and nonlinear intensity adjustment to approximate BOLD contrast while preserving anatomical geometry. BOLD references un-derwent sequential rigid, affine, and nonlinear SyN registration to these targets, followed by a second synthetictarget estimation and SyN refinement [100]. Transformations were combined and applied to the full BOLD time series independently for LR and RL acquisitions. Corrected BOLD data were coregistered to each animal’s anatomical image (ANTs, rigid-body) and warped to the NMT v2.0 template via the subject-specific anatomical template.

Denoising consisted of nuisance regression with 24 head-motion regressors (six rigid-body estimates, their temporal derivatives, and the squares of both)[101], the first five anatomical CompCor components derived from white-matter and cerebrospinal-fluid masks, and the global signal [102]. Linear detrending, band-pass filtering (0.01Hz-0.08Hz), and spatial smoothing (3mm FWHM Gaussian kernel) were additionally performed.

Functional connectomes were constructed using the CHARM level-6 cortical parcellation and selected SARM subcortical regions spanning the midbrain, brainstem, and cerebellum (levels 2, 3, and 5) [87, 94]. The combined atlas contained 316 nodes—264 cortical and 52 subcortical—after excluding four cortical regions with insufficient coverage at the functional resolution (left and right posterior olfactory cortex [OT/Pir] and left and right area TH). The two denoised phase-encoding runs were concatenated per session, and mean time series were extracted from each node, forming the basis of the connectivity analyses described below.

In addition to the anatomical networks defined by the CHARM level-1 labels, cortical parcels were classified into the seven canonical functional networks defined by Yeo et al. [103] using the human-to-macaque functional correspondence established by Xu et al. [104]. The network labels were transformed into NMT v2.0 space using RheMAP, and each parcel’s assignment was verified against the modal voxelwise network label within its boundaries in the NMT v2.0 implementation of the Yeo atlas. Hemisphere was determined from the RAS+ world-coordinate centroid of each parcel.

### Covariance estimation and Riemannian centering

Longitudinal functional-connectivity matrices carry stable, individual-specific signatures that can dominate smaller state-dependent differences [43, 44, 105, 106]. To separate these stable between-animal offsets from the within-animal variation of interest, we represented each scan as a regional covariance matrix and aligned the animal-specific means in Riemannian space.

For each animal and session (13 × 6 = 78 scans), we estimated a 316 × 316 region-wise covariance matrix from the denoised regional time series. To ensure numerical positive-definiteness, we applied 1% tracepreserving shrinkage toward the identity:

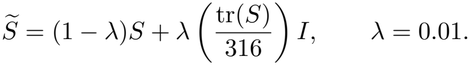

We retained covariance rather than rescaling the matrices to correlation because covariance preserves regional differences in BOLD signal variance within the symmetric positive-definite representation used for centering.

Riemannian centering followed [45], and as implemented in our previous work [34, 35, 37, 46, 107, 108]. The procedure maps each animal’s geometric-mean co-variance to the grand geometric mean while preserving the relative structure of its session-level matrices. We first computed the grand geometric mean, S_*gm*_, across all 78 covariance matrices. For each animal *i*, we computed the geometric mean of its six session-specific covariance matrices, S_*i*_, and mapped each session matrix, S_*ij*_, onto the tangent space at S_*i*_:

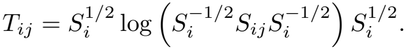

Each tangent vector was then transported to the tangent space at the grand mean:

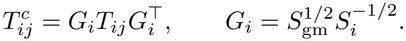

Finally, the transported tangent vector was mapped back onto the manifold of symmetric positive-definite co-variance matrices:

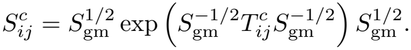

Here, log and exp denote the matrix logarithm and matrix exponential, respectively.

### Whole-connectome multivariate distance matrix regression

Before localizing functional reorganization to individual regions, we first asked, as an omnibus test, whether housing condition altered the connectome as a whole. We tested this using multivariate distance matrix regression (MDMR), a method that relates pairwise differences among complete connectivity matrices to explanatory variables [47–49]. Pairwise distances among the 78 centered covariance matrices were calculated as one minus the Pearson correlation between their vectorized upper-triangular edges. The model included housing condition and centered linear session, corresponding to the fixed effects in the common longitudinal model, together with subject as a fixed blocking factor, which removes each animal’s mean connectome in place of the random intercept. The housing-condition effect was evaluated with linear session held constant using a pseudo-*F* statistic, with effect size expressed as partial *R*2; the session effect was evaluated analogously with housing condition held constant.

Because each animal contributed six connectivity matrices, statistical significance could not be assessed by unrestricted permutation across all 78 scans. Housing-condition labels were shuffled among the six sessions within each animal, with the subject blocking factor and linear session covariate kept fixed. Permutation-based *p* values were estimated from 5,000 permutations.

### Functional-connectivity manifold analysis

#### Manifold construction

To represent each region according to its pattern of connectivity with the rest of the brain, we constructed a low-dimensional manifold from each centered covariance matrix. This approach followed established connectivity-gradient methods [32, 38, 50] and our previous applications of manifold learning to task- and state-dependent brain reorganization in humans and nonhuman primates [34, 35, 109, 110].

Within each matrix, connections were thresholded row-wise to retain the strongest 10% of positive values for each region. Because thresholding was performed independently for each row, the resulting matrix could be asymmetric and was not symmetrized. We then calculated the cosine similarity between every pair of regional connectivity profiles, producing a symmetric affinity matrix in which larger values indicated greater similarity in how two regions connected with the rest of the brain.

Each affinity matrix was decomposed using diffusion map embedding [111], as implemented in BrainSpace [112] (anisotropic diffusion parameter *α* = 0.5; diffusion time t = 0). This procedure assigned each region coordinates along continuous gradients capturing the principal dimensions of variation among whole-brain connectivity profiles. Ten gradients were computed and the first three were retained for the primary analyses. Repeating the analyses with more than three gradients yielded highly similar patterns of regional effects (see *Robustness and sensitivity analyses*). The relative contribution of each gradient was expressed as its eigenvalue divided by the sum of all ten computed diffusion-map eigenvalues.

#### Template alignment

To compare regional manifold coordinates across animals and sessions, the separately estimated embeddings were expressed in a common co-ordinate system; without alignment, equivalent gradients can differ in sign and orientation. We constructed a template manifold by applying the same embedding procedure to the mean of the centered covariance matrices across all 78 scans. Each scan-specific manifold was then aligned to this template using an orthogonal Procrustes transformation [113], applied without centering or scaling. Because the transformation is orthogonal, it preserves each region’s distance from the manifold centroid and therefore leaves eccentricity unchanged. Repeating alignment with a template derived from the first-session matrices alone yielded near-identical regional eccentricity statistics, and the corresponding template component maps were highly similar (fig. S6B-D). Template choice affects the orientation used to display and compare gradient coordinates, rather than the eccentricity measure itself.

#### Manifold eccentricity

We summarized each region’s position within the connectivity manifold using manifold eccentricity, which is the Euclidean distance between its coordinates across the first three gradients and the manifold centroid [33, 34, 110]. For region *i* in animal *j* at session *t*:

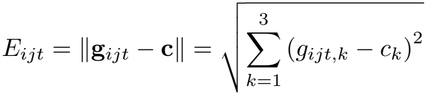

Where

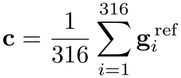

Here, **g**_*ijt*_ is the vector containing the region’s coordinates along the three aligned gradients, and **c** is the manifold centroid — the mean position of all 316 regions within the template embedding, a single fixed anchor shared across all animals and sessions. Larger values indicate that a region’s whole-brain connectivity profile is more strongly differentiated along one or more of the principal gradients, whereas smaller values indicate that its profile lies closer to the centroid and is less strongly expressed along those dimensions. Eccentricity is therefore a geometric summary of connectivity differentiation, rather than a direct measure of mean connectivity strength.

We calculated eccentricity for each of the 316 regions in every animal and session. Increases in eccentricity across housing phases were termed manifold expansion, and decreases were termed manifold contraction.

#### Statistical inference on eccentricity

Condition-related changes in eccentricity were tested separately for each of the 316 regions using the common longitudinal model defined above, with eccentricity as the outcome. Models were fitted using Python’s statsmodels v0.14.6, and the housing-condition term was evaluated using Wald *z* statistics. Statistical significance was determined by applying Benjamini–Hochberg false-discovery-rate correction across all 316 regions (*q* < 0.05 ) [114].

#### Robustness and sensitivity analyses

We examined sensitivity of the eccentricity findings to BOLD signal variance, model specification, manifold geometry and the principal analytical choices. Sensitivity was assessed through the direction and spatial pattern of effects as well as the number of regions surviving multiplicity correction; these summaries distinguish agreement in effect estimates from changes in statistical precision. The out-comes of these analyses are reported in the Results and figs. S5-7.

Because covariance preserves regional differences in signal amplitude, we tested whether BOLD variance could account for the eccentricity effects (fig. S5). First, we fit the longitudinal model to each region’s BOLD variance to test whether amplitude itself changed with housing. Second, for each region we quantified the withinregion association between eccentricity and BOLD variance as the Pearson correlation across the 78 scans. Third, we refit the eccentricity model with each region’s BOLD variance added as a covariate, testing whether the condition effect survived adjustment for amplitude.

We next assessed sensitivity to model specification by replacing the random-intercept model with a repeated-measures analysis of variance containing fixed animal effects. As a complementary anatomical-plausibility check, we compared condition-effect estimates between left– right homologous regions matched by atlas label. This comparison was treated as a measure of bilateral concordance rather than as an independent replication.

Finally, we repeated the complete manifold pipeline across its principal analytical choices: row-retention thresholds from 5% to 100% in 5% steps; one to ten retained gradients; and an alternative alignment template (the first-session mean in place of the grand mean). The outcomes of these sensitivity analyses are reported in fig. S6.

#### Connectivity changes associated with eccentricity effects

To describe the connectivity changes accompanying the regional eccentricity effects, we derived each region’s connectivity-change profiles. That is, its change in Fisher *z* connectivity to every other region between conditions (the mean difference between paired housing sessions and single housing sessions). Because these profiles were from regions selected for their eccentricity effects in the same data, they were used to characterize the detected regional effects, not to make independent edge-level claims. Representative profiles are shown in Fig 4A.

To identify shared patterns of reorganization among the condition-responsive regions, each region’s connectivity-change profile was treated as a feature vector, and pairwise dissimilarity was computed as the correlation distance (1 - *r*). The dissimilarity matrix was visualized with metric multidimensional scaling (MDS). Regions were then grouped by agglomerative hierarchical clustering with average linkage on the same correlation-distance matrix, and the number of clusters was selected over k = 2-10 using the silhouette coefficient [115] and the Calinski–Harabasz criterion [116], which supported a two-cluster solution.

Connectivity changes underlying each mode were summarized at the network level. For each significant region, its connectivity change profile was averaged across the connections to each target network (anatomical divisions and functional networks); regions were grouped by mode (Fig. 4E). The correspondence between the two modes was quantified as the correlation between their network-level means.

Mode-level connectivity patterns were examined by taking the mean Fisher *z* value within each mode (within-mode connectivity) and between both modes (between-mode connectivity) for each scan. The segregation index was additionally computed based on these values [117]. For each animal, these measures were averaged within the single- and paired-housing conditions and used for descriptive analyses between conditions.

These analyses were restricted to regions identified by the eccentricity analysis and were therefore intended to characterize the detected responses, not to provide an independent test of the housing-condition effect. Because clustering was based on condition-related connectivity changes rather than baseline connectivity, the resulting groups represent shared modes of reorganization across the housing transition rather than conventional resting-state networks.

### Hair cortisol collection, assay and analysis

Hair was collected approximately one week before each imaging session by shaving the lumbar region. At the initial collection, following approximately nine months of stable single housing, only the proximal half of the hair shafts—the half closest to the skin—was analyzed to favor more recently formed hair. This baseline sample’s integration period was not defined by a preceding shave. At subsequent collections, the same region was re-shaved, so samples from sessions 2-6 comprised hair grown since the preceding collection and reflected cumulative cortisol exposure over approximately three months, consistent with established hair-sampling methods in macaques [39]. Samples were sealed in aluminum foil and stored dry at ^−^ 80 °C until processing. Of the 78 scheduled collections, 75 yielded sufficient hair for analysis, with sample sizes of 13, 13, 13, 11, 12, and 13 at sessions 1-6, respectively.

Hair cortisol was quantified by the Drug Safety Laboratory at the Robarts Research Institute, Western University, using its established hair-analysis procedure [118, 119]. Briefly, samples were weighed, washed with iso-propanol to remove external contamination, air-dried, pulverized, and extracted with an organic solvent. The supernatant was evaporated to dryness under nitrogen, and the residue was reconstituted in phosphate-buffered saline before cortisol measurement. Hair cortisol concentrations were analyzed in their original laboratory-reported units (ng/g), without further transformation or normalization, using the common longitudinal model defined above. Missing samples were not imputed. Because the session 1 sample was not preceded by a shave and was restricted to the proximal half of the hair shaft, we also fitted the same model to sessions 2–6 only (62 samples from 13 animals). This analysis showed that, even without session 1, the paired-housing effect was present (*β* = 56.3 ng/g, 95% CI [26.2, 86.4], Wald *z* = 3.67, *p* < 0.001).

### General statistical procedures

All statistical tests were two-sided and used an alpha level of 0.05. Effect estimates and 95% confidence intervals are reported alongside *P* values, and multiplicity correction was applied as described for each analysis. Each animal contributed repeated measurements, and housing partners shared one of six social units. The fixed A-B-A housing sequence was not randomized, and animal-care personnel could not be blinded to housing condition. Partner assignment was based on veterinary and behavioral compatibility rather than random allocation, as described under Experimental design.

## Data and code availability

The derived data and analysis materials required to reproduce the reported findings will be deposited in Github upon publication. These materials will include deidentified regional measurements, centered connectome matrices, model specifications, analysis code, software-environment information, quality-control summaries, complete statistical outputs, and the source data underlying the figures and tables. Raw data are available on request by contacting the senior author and under an inter-institutional Data Transfer Agreement with Queen’s University. Source data are provided with this paper.

## Acknowledgements

We would like to give a special thanks to the Animal Care Staff at Queen’s University for making this work possible, specifically Brittney Armitage-Brown, Kim Moore, Jessica Freeman, Angie Thompson, Darlene Potterton, and Kurt Scrutton, for their assistance with this work. Funding for the project was provided by the New Frontiers in Research Fund (NFRFE-2021-00936) awarded to J.P.G and A.T., an Ontario Early Researcher Award (ER19-15-156) awarded to J.P.G., and internal funding provided by Queen’s University through the Harry Botterell Foundation for the Neurological Sciences and the Vice-Principal Research portfolio.

## Author contributions

Conceptualization: S.B., A.W. & J.P.G. Methodology: D.J.G., J.Y.N., C.P., D.B. & J.P.G. Data Acquisition: D.J.G., J.Y.N., S.B., A.W. & J.P.G. Formal analysis: D.J.G., J.Y.N., C.P., & J.P.G. Software: D.J.G. Validation: D.J.G. & J.P.G. Data curation: D.J.G., J.Y.N., C.P., S.B., A.W., & J.P.G. Visualization: D.J.G & J.P.G. Resources: A.R., S.L.R., M.A.E., C.A., I. G-H., D.B., J.S., V. A.K., M.S., E.R.O., P.M.S., G.B., M.P., S.S., D.P.M., F.G.D., D.J.C., S.B., A.W. & J.P.G. Funding acquisition: A.T., D.J.C., S.B., A.W. & J.P.G. Project administration: S.B., A.W., & J.P.G. Supervision: S.B., A.W., & J.P.G. Writing—original draft: D.J.G. & J.P.G. Writing—review & editing: D.J.G., J.Y.N., C.P., A.R., S.L.R., M.A.E., I.G-H., D.B., V. A.K., M.S., J.D.W., E.R.O., P.M.S., G.B., M.P., S.S., D.P.M., F.G.D., D.J.C., A.T., S.B., A.W. & J.P.G. All authors approved the final manuscript.

## Notes

### Competing Interest Statement

The authors have declared no competing interest.

