## Supplemental Materials for "Social environment induces reversible remodeling of the adult primate brain"

**Supplemental Figure 1:** Mid-STS expansion converges anatomically with social-network effects in previous macaque studies

**Supplemental Figure 2:** Reduced statistical thresholds reveal bilateral expression of the structural housing effects

**Supplemental Figure 3:** Grey-matter changes consistently reverse direction after paired housing ends

**Supplemental Figure 4:** Phase-specific contrasts localize the emergence and reversal of grey-matter expansion

**Supplemental Figure 5:** Paired-housing eccentricity effects persist after accounting for regional BOLD variance

**Supplemental Figure 6:** The regional eccentricity pattern is retained across model and embedding choices

**Supplemental Figure 7:** Unthresholded paired-housing eccentricity effects and bilateral correspondence

**Supplemental Figure 8:** Hair cortisol concentrations are higher during paired housing

**Supplemental Table 1:** Grey-matter expansion during protected-contact paired housing

**Supplemental Table 2:** Regions with significant paired-housing eccentricity effects

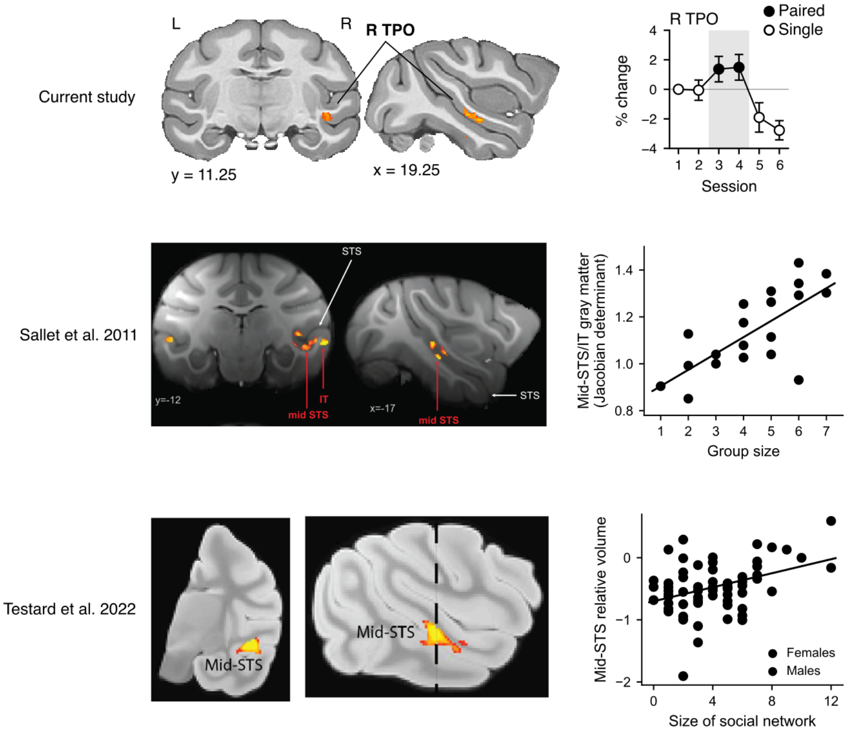


**Fig.** S1. **Mid-STS expansion converges anatomically with social-network effects in previous macaque studies |** Top, anatomical location and session-resolved grey-matter change for the right TPO cluster in the current study, showing expansion during paired housing and reversion after single housing resumed. Middle, mid-STS cluster identified by Sallet et al. [1], with greater local grey-matter volume in macaques housed in larger social groups. Bottom, mid-STS cluster from Testard et al. [2], with greater relative volume in free-ranging macaques with larger social networks ($r$ = 0.38, $p$ = 0.001); purple and green symbols denote females and males, respectively. This anatomical convergence across experimental and free-ranging settings links the present reversible response to a reproducible locus of social-environment-related structural variation. Scatterplots were digitized from Sallet et al., Fig. 1A (20 of 23 animals recovered; three markers fully occluded), and Testard et al., Fig. 1 (61 of approximately 68 animals recovered), and replotted in a common format. Black lines reproduce the published regression fits. Brain image adapted from Testard et al. (2022), under a CC BY 4.0 licence.

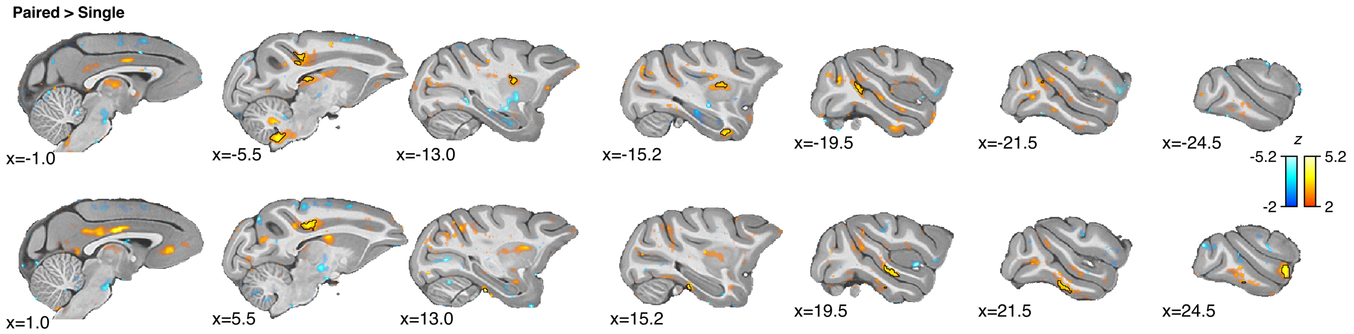


**Fig.** S2. **Reduced statistical thresholds reveal bilateral expression of the structural housing effects |** Sagittal sections show paired-versus-single housing differences at an uncorrected voxelwise $p < 0.05$ (two-sided; displayed $|z| \geq 2$). Note that many effects extend across corresponding territories in both hemispheres, including temporal and medial cortical regions, supporting the spatial consistency of the housing-related pattern beyond the clusters surviving the principal corrected analysis in Fig. 2. Warm and cool colors indicate greater and lower grey-matter volume during pairing, respectively. Black outlines delineate the significant clusters identified in the principal analysis (voxelwise $p < 0.001$; cluster-corrected $p < 0.05$). Slice positions are given in millimeters in NMT v2 space.

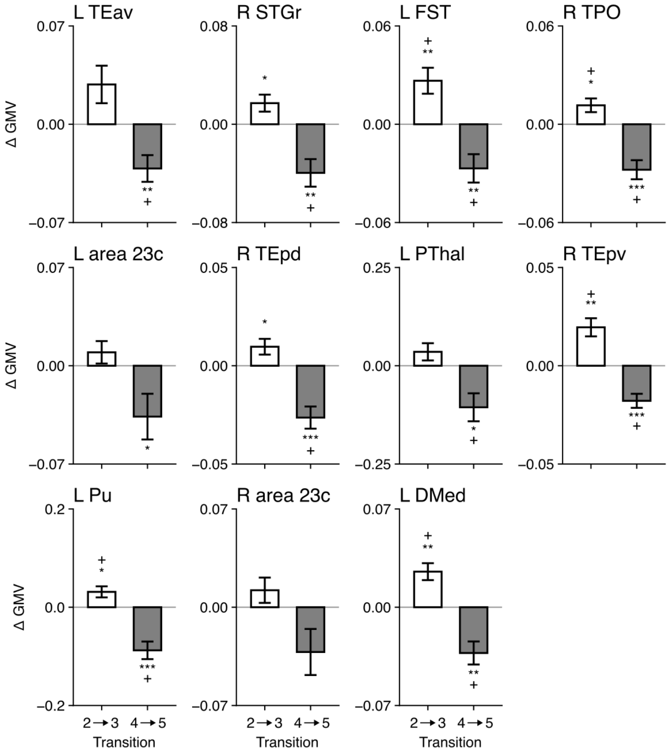


**Fig.** S3. **Grey-matter changes consistently increase during paired housing and then reverse direction after paired housing ends |** Mean within-animal change in grey-matter volume across the two housing transitions, for each of the 11 clusters identified by the principal paired-versus-single contrast (Fig. 2; cluster abbreviations follow Table S1). White bars show the change after pairing began (session 3 minus session 2) and grey bars show the change after single housing resumed (session 5 minus session 4). ΔGMV is the between-session difference in each animal's mean modulated grey-matter value within the cluster. Bars show the mean across the 13 macaques and error bars the SEM of the within-animal differences. Asterisks mark uncorrected two-sided paired *t*-tests (*$p < 0.05$, **$p < 0.01$, ***$p < 0.001$), and + marks changes that also pass Benjamini-Hochberg correction across the 11 clusters ($q < 0.05$). These markers are descriptive. The clusters were selected using the paired-versus-single contrast, which is computed from the same sessions, so the tests are not independent of cluster selection and do not provide separate evidence for emergence or reversal. Whole-brain tests of the phase contrasts are shown in Fig. S4.

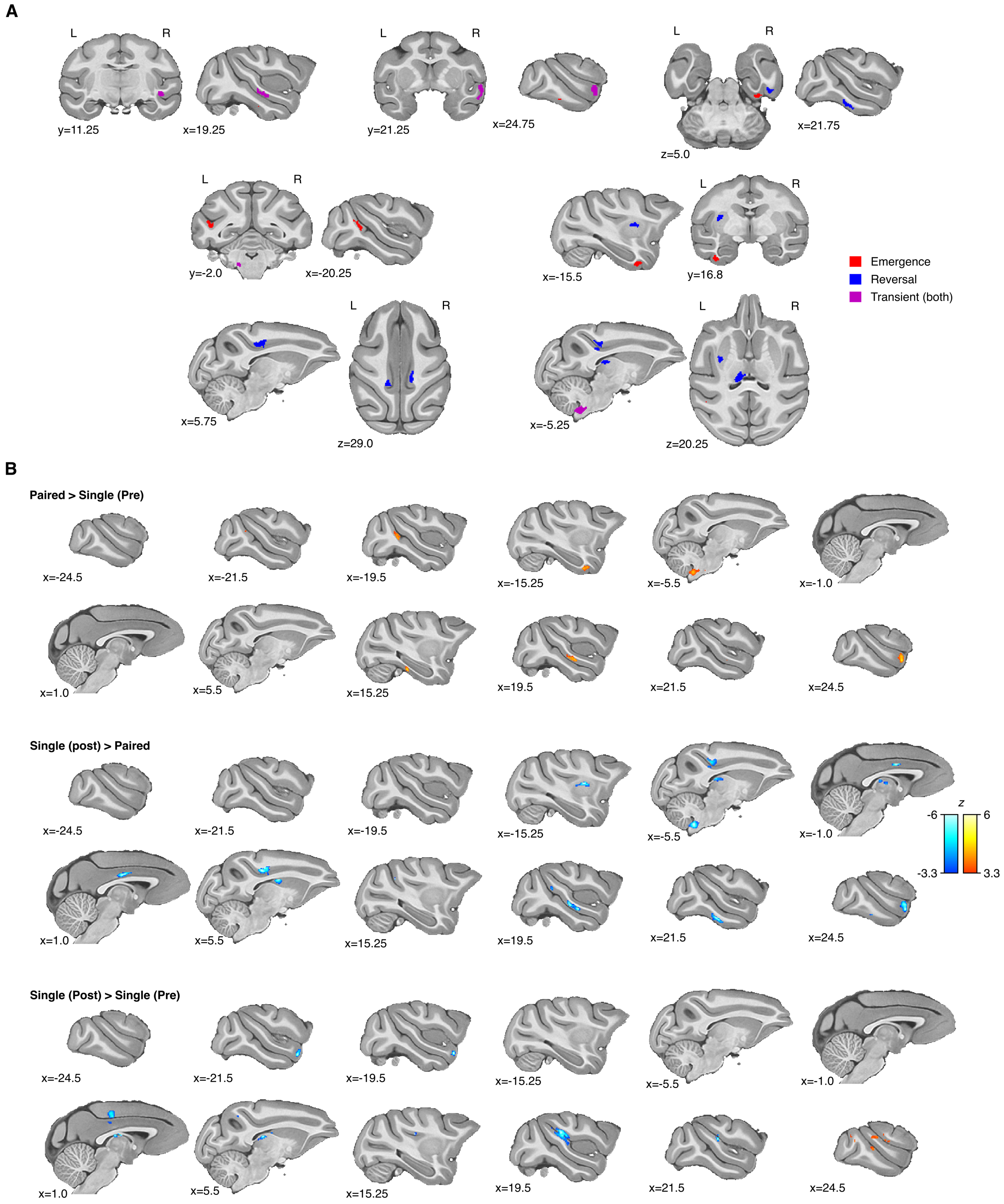


**Fig.** S4. **Phase-specific contrasts localize the emergence and reversal of grey-matter expansion |** Single housing was separated into Pre (sessions 1-2) and Post (sessions 5-6), with Paired comprising sessions 3-4. (A) Spatial correspondence of the 11 principal clusters with separate whole-brain phase contrasts: three clusters overlapped the emergence contrast only (Paired > Pre; red), five the reversal contrast only (Post > Paired; blue), and three for both contrasts (magenta). None overlapped the Post > Pre contrast. Overlap was calculated as the fraction of a cluster's voxels within each contrast's separately thresholded maps. Overlaps ranged between 76-99%. (B) Whole-brain maps for Paired > Pre (top), Post > Paired (middle), and Post > Pre (bottom), each thresholded at voxelwise $p < 0.001$ and cluster-corrected $p < 0.05$. Positive values in the top maps indicate expansion during pairing; negative values in the middle maps indicate recession after pairing ended. These spatially convergent contrasts characterize the emergence and reversal of the structural response across the same longitudinal scans.

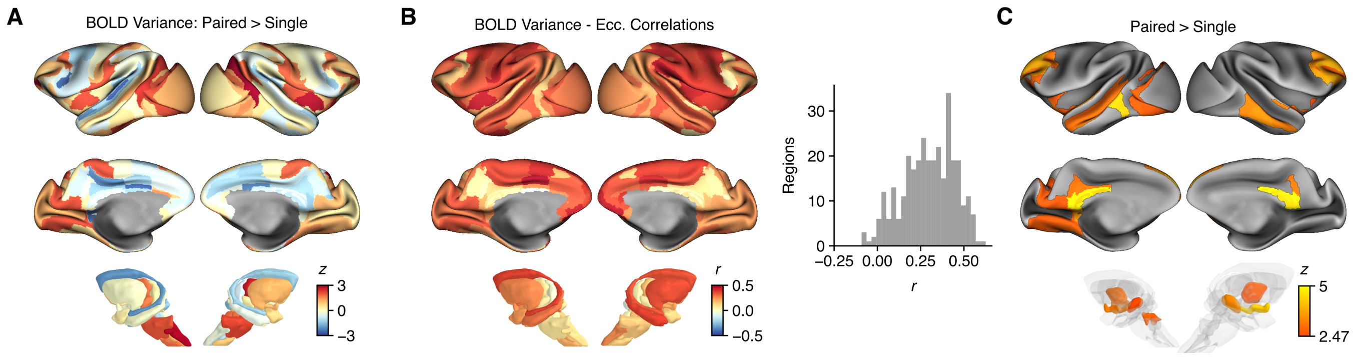


**Fig.** S5. **Paired-housing eccentricity effects persist after accounting for regional BOLD variance |** (A) Unthresholded condition-effect $z$ statistics for regional BOLD variance from the longitudinal model; none of the 316 regions survived FDR correction ($q < 0.05$). (B) Regional BOLD variance and manifold eccentricity are predominantly positively associated across scans, shown as a correlation map and r-distribution across regions. (C) Eccentricity condition effects after adding each region's BOLD variance to the longitudinal model. 80 of the 88 regions identified in the principal analysis remained significant (FDR $q < 0.05$), with positive $z$ values indicating greater eccentricity during pairing. Thus, regional signal variance covaries with eccentricity but does not account for the majority of its housing-related effects.

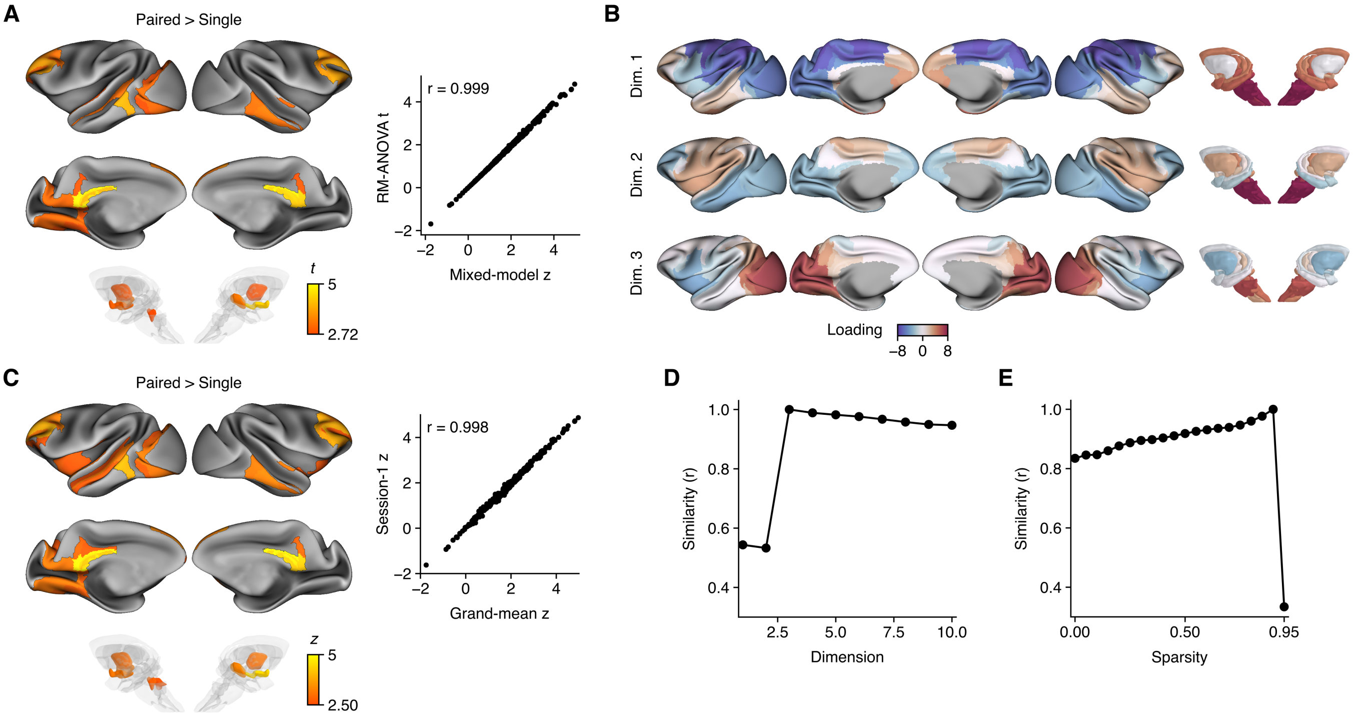


**Fig.** S6. **The regional eccentricity pattern is retained across model and embedding choices |** (A) A repeated-measures ANOVA with fixed animal effects yields condition statistics nearly identical to the mixed-effects model ($r = 0.999$ across regions); 58 of the 88 principal regions remain significant (FDR $q < 0.05$). Positive condition statistics indicate greater eccentricity during pairing. (B) The first three dimensions of a template manifold constructed from the mean connectivity across session 1 scans only, illustrating a highly similar large-scale organization at study baseline. (C) Regions with significant condition effects from the mixed-effects model when eccentricity is derived from the session-1 template manifold in (B) rather than the grand-mean template; 85 of the 88 regions remain significant. Wald $z$ shown thresholded at FDR $q < 0.05$. (D and E) Spatial correlations of regional condition statistics with the primary analysis (three gradients; strongest 10% of connections retained per row), varying the number of gradients (D) or sparsity (E). Agreement remains high with 3–10 gradients ($r \geq 0.95$) but falls when only one or two gradients are retained ($r = 0.53$-$0.54$). Across sparsity settings, correspondence was high ($r \geq 0.8$) and peaked at the primary setting (0.9), except at the highest sparsity of 0.95, which collapsed to $r = 0.33$.

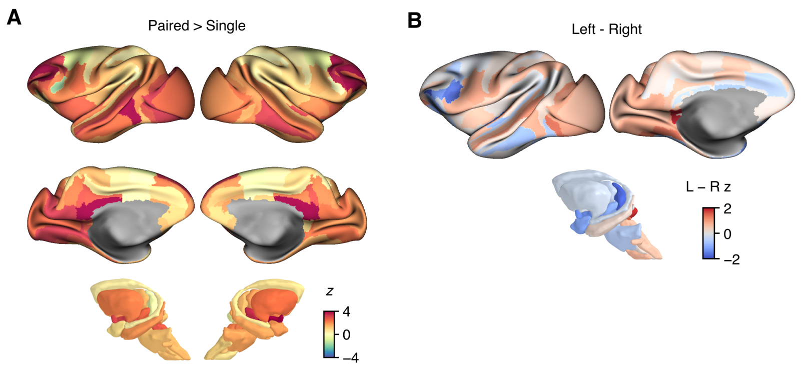


**Fig.** S7. **Unthresholded paired-housing eccentricity effects and bilateral correspondence |** (A) Unthresholded paired-versus-single condition-effect Wald $z$ statistics show corresponding patterns across hemispheres; positive values denote greater eccentricity during pairing. Statistics in left–right homologues, matched by atlas label, are strongly correlated ($r = 0.79$). (B) Left-minus-right differences in these statistics illustrate the regional departures from this overall bilateral correspondence.

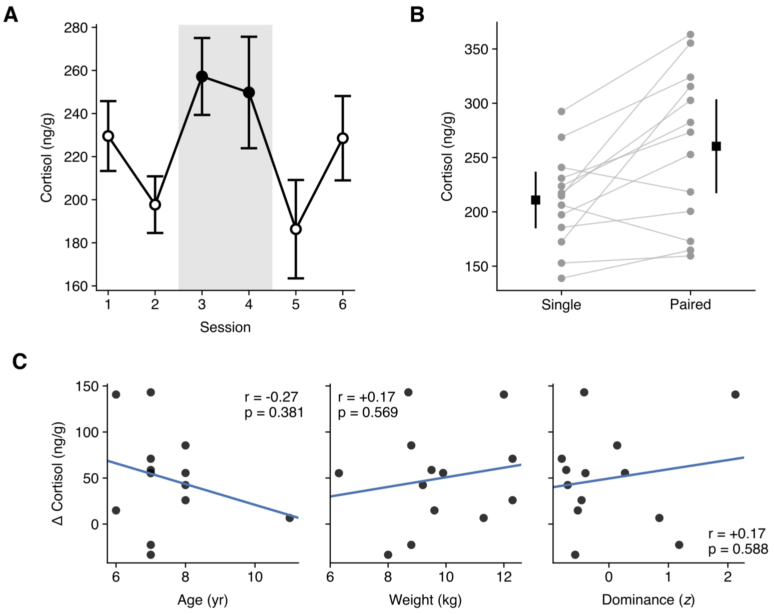


**Fig.** S8. **Hair cortisol concentrations are higher during paired housing |** (A) Session-resolved hair cortisol concentrations across the single–paired–single housing protocol, also shown in Fig. 1C. Points show the mean ± SEM. Shading and filled symbols indicate paired housing (sessions 3–4); open symbols indicate single housing. The longitudinal model identified higher cortisol concentrations during paired housing ($\beta$ = 45.9 ng/g, 95% CI 19.2–72.6, $p$ = 0.0008). The estimate was similar when session 1 was excluded ($\beta$ = 56.3 ng/g, 95% CI 26.2–86.4, $p$ = 0.0002). (B) Within-animal comparison of mean cortisol concentrations during single housing (sessions 1, 2, 5 and 6) and paired housing (sessions 3–4). Grey lines connect individual animals; black squares and whiskers show the group mean ± 95% CI. Concentrations were higher during paired housing in 11 of 13 animals (mean paired-minus-single difference = 49.5 ng/g, 95% CI 17.0–82.0; paired $t(12)$ = 3.32, $p$ = 0.006). (C) Pairing-related cortisol changes (paired minus single mean) plotted against age, body weight and dominance (mean Dominant rating, standardized within each phase and averaged across phases). Lines show least-squares fits; $r$ and $p$ values indicate Pearson correlations (n = 13; uncorrected). No statistically significant associations were detected. These exploratory correlations do not exclude a relationship between observer-rated dominance and cortisol change.

**Table** S1. **Grey-matter expansion during protected-contact paired housing |** Eleven clusters showed greater grey-matter volume during paired than single housing in the longitudinal model (voxelwise $p < 0.001$; cluster-corrected $p < 0.05$). Coordinates (x, y, z) are in millimeters in NMT v2 space; peak Wald $z$ statistics and cluster volumes ($mm^3$) are reported.

| Cluster | Region | x (mm) | y (mm) | z (mm) | Peak z | Volume ($mm^3$) |
|---|---|---|---|---|---|---|
| 1 | R TEpv | 16.2 | 3.4 | 4.6 | 5.42 | 17.5 |
| 2 | L DMed | -6.2 | -4.1 | -0.9 | 5.38 | 31.4 |
| 3 | R STGr | 24.8 | 21.9 | 9.6 | 5.25 | 27.6 |
| 4 | R 23c | 6.2 | 6.9 | 28.1 | 5.19 | 21.6 |
| 5 | R TEpd | 23.8 | 5.4 | 6.1 | 4.72 | 23.8 |
| 6 | R TPO | 19.8 | 12.9 | 9.6 | 4.7 | 25.6 |
| 7 | L TEav | -16.2 | 16.4 | -0.4 | 4.67 | 17.5 |
| 8 | L PThal | -5.2 | 6.4 | 20.6 | 4.62 | 22.9 |
| 9 | L Pu | -13.8 | 15.9 | 19.6 | 4.55 | 25.5 |
| 10 | L 23c | -4.8 | 3.4 | 26.6 | 4.46 | 20.9 |
| 11 | L FST | -19.8 | -1.1 | 15.6 | 4.39 | 18.2 |

**Table** S2. **Regions with significant paired-housing eccentricity effects from the mixed-effects model |** For each of the 88 regions, hemisphere, anatomical and functional (Yeo 7-network) network assignments, Wald $z$, and uncorrected $p$. All regions listed survived Benjamini–Hochberg FDR correction ($q < 0.05$ across 316 regions).

| CHARM/SARM region | Hemi. | Anatomical network | Yeo network | Wald $z$ | $p$ |
|---|---|---|---|---|---|
| area_v23 | L | Parietal | Default mode | 4.97 | <0.001 |
| area_23a | L | Parietal | Default mode | 4.82 | <0.001 |
| area_23a | R | Parietal | Default mode | 4.81 | <0.001 |
| area_29 | R | Parietal | Frontoparietal | 4.52 | <0.001 |
| area_30 | L | Parietal | Default mode | 4.47 | <0.001 |
| subpallial_amygdala | R | Limbic | Subcortical | 4.35 | <0.001 |
| area_v23 | R | Parietal | Default mode | 4.32 | <0.001 |
| area_30 | R | Parietal | Default mode | 4.28 | <0.001 |
| area_29 | L | Parietal | Frontoparietal | 4.11 | <0.001 |
| area_45a | R | Frontal | Default mode | 3.94 | <0.001 |
| area_TEO | L | Temporal | Visual | 3.92 | <0.001 |
| rostromedial_belt_region | R | Temporal | Limbic | 3.89 | <0.001 |
| dorsal_area_46 | R | Frontal | Default mode | 3.89 | <0.001 |
| dorsal_area_46 | L | Frontal | Ventral att. | 3.88 | <0.001 |
| rostral_area_12 | R | Frontal | Default mode | 3.84 | <0.001 |
| ventral_area_46 | L | Frontal | Default mode | 3.83 | <0.001 |
| arcuate_sulcus_area_8B | R | Frontal | Frontoparietal | 3.82 | <0.001 |
| floor_of_the_superior_temporal_area | L | Temporal | Dorsal att. | 3.82 | <0.001 |
| ventral_area_46 | R | Frontal | Default mode | 3.8 | <0.001 |
| area_45b | R | Frontal | Frontoparietal | 3.71 | <0.001 |
| medial_area_12 | R | Frontal | Default mode | 3.69 | <0.001 |
| rostrotemporal_core_region | R | Temporal | Limbic | 3.54 | <0.001 |
| rostral_dorsal_premotor_cortex | R | Frontal | Default mode | 3.53 | <0.001 |
| rostral_dorsal_premotor_cortex | L | Frontal | Default mode | 3.48 | <0.001 |
| dorsal_area_9 | L | Frontal | Default mode | 3.46 | <0.001 |
| geniculate_thalamus | R | Thalamus | Subcortical | 3.4 | <0.001 |
| dorsal_periarcuate_area_8A | R | Frontal | Frontoparietal | 3.39 | <0.001 |
| polar_rostrotemporal_cortex | R | Temporal | Limbic | 3.33 | <0.001 |
| polar_rostrotemporal_cortex | L | Temporal | Limbic | 3.32 | <0.001 |

**Table** S2. (continued)

| | | | | | |
|---|---|---|---|---|---|
| dorsal_area_8B | L | Frontal | Default mode | 3.3 | <0.001 |
| lateral_rostrotemporal_belt_region | L | Temporal | Limbic | 3.3 | <0.001 |
| rostrotemporal_core_region | L | Temporal | Limbic | 3.29 | <0.001 |
| dorsal_area_8B | R | Frontal | Default mode | 3.27 | 0.001 |
| hypothalamus | L | Limbic | Subcortical | 3.25 | 0.001 |
| dorsal_area_9 | R | Frontal | Default mode | 3.25 | 0.001 |
| clearly_part_of_visual_area_2 | L | Occipital | Visual | 3.23 | 0.001 |
| posterior_dorsal_area_TE | R | Temporal | Limbic | 3.23 | 0.001 |
| area_PGa | L | Temporal | Default mode | 3.2 | 0.001 |
| arcuate_sulcus_area_8B | L | Frontal | Frontoparietal | 3.18 | 0.001 |
| anterior_dorsal_area_TE | R | Temporal | Limbic | 3.12 | 0.002 |
| area_TEO | R | Temporal | Visual | 3.11 | 0.002 |
| rostral_parabelt_region | R | Temporal | Default mode | 3.1 | 0.002 |
| area_TEa | R | Temporal | Limbic | 3.1 | 0.002 |
| subpallial_amygdala | L | Limbic | Subcortical | 3.04 | 0.002 |
| area_46_in_fundus_of_the_principal_sulcus | L | Frontal | Frontoparietal | 3.02 | 0.002 |
| globus_pallidus | L | Basal ganglia | Subcortical | 2.94 | 0.003 |
| anterior_lateral_belt_region | L | Temporal | Somatomotor | 2.9 | 0.004 |
| dorsal_visual_area_3 | L | Occipital | Visual | 2.89 | 0.004 |
| medial_agranular_insular_region | R | Frontal | Limbic | 2.89 | 0.004 |
| dorsal_periarcuate_area_8A | L | Frontal | Frontoparietal | 2.86 | 0.004 |
| globus_pallidus | R | Basal ganglia | Subcortical | 2.86 | 0.004 |
| floor_of_the_superior_temporal_area | R | Temporal | Ventral att. | 2.85 | 0.004 |
| area_31 | R | Parietal | Somatomotor | 2.85 | 0.004 |
| ventral_visual_area_3 | L | Occipital | Visual | 2.84 | 0.005 |
| area_31 | L | Parietal | Somatomotor | 2.83 | 0.005 |
| lateral_pons | L | Brainstem/Mid. | Subcortical | 2.79 | 0.005 |
| possibly_part_of_visual_area_2 | L | Occipital | Visual | 2.77 | 0.006 |
| lateral_area_11 | L | Frontal | Limbic | 2.76 | 0.006 |
| lateral_rostrotemporal_belt_region | R | Temporal | Limbic | 2.75 | 0.006 |

**Table** S2. (continued)

| | | | | | |
|---|---|---|---|---|---|
| lateral_agranular_insula_area | L | Frontal | Default mode | 2.75 | 0.006 |
| posterior_lateral_agranular_insula_area | L | Frontal | Default mode | 2.71 | 0.007 |
| visual_area_V3A | L | Occipital | Visual | 2.71 | 0.007 |
| area_23b | L | Parietal | Ventral att. | 2.7 | 0.007 |
| medial_area_12 | L | Frontal | Default mode | 2.69 | 0.007 |
| area_45a | L | Frontal | Default mode | 2.68 | 0.007 |
| gustatory_cortex | L | Frontal | Ventral att. | 2.67 | 0.008 |
| parainsula | L | Temporal | Limbic | 2.67 | 0.008 |
| area_TEa | L | Temporal | Limbic | 2.64 | 0.008 |
| dorsal_pons | L | Brainstem/Mid. | Subcortical | 2.64 | 0.008 |
| posterior_lateral_agranular_insula_area | R | Frontal | Frontoparietal | 2.64 | 0.008 |
| lateral_pallium | R | Limbic | Subcortical | 2.63 | 0.008 |
| parainsula | R | Temporal | Limbic | 2.63 | 0.009 |
| medial_rostrotemporal_belt_region | L | Temporal | Limbic | 2.62 | 0.009 |
| agranular_and_dysgranular_insula | L | Temporal | Ventral att. | 2.61 | 0.009 |
| medial_rostrotemporal_belt_region | R | Temporal | Limbic | 2.6 | 0.009 |
| ventral_periarcuate_area_8A | R | Frontal | Frontoparietal | 2.6 | 0.009 |
| temporal_parietooccipital_associated_area | L | Temporal | Default mode | 2.59 | 0.010 |
| area_IPa | L | Temporal | Default mode | 2.58 | 0.010 |
| area_46_in_fundus_of_the_principal_sulcus | R | Frontal | Frontoparietal | 2.58 | 0.010 |
| orbital_frontal_pole | L | Frontal | Default mode | 2.57 | 0.010 |
| lateral_agranular_insula_area | R | Frontal | Default mode | 2.56 | 0.011 |
| rostral_core_region | L | Temporal | Somatomotor | 2.55 | 0.011 |
| lateral_pallium | L | Limbic | Subcortical | 2.52 | 0.012 |
| secondary_somatosensory_cortex | L | Parietal | Ventral att. | 2.5 | 0.012 |
| primary_visual_cortex | L | Occipital | Visual | 2.49 | 0.013 |
| area_TEm | L | Temporal | Limbic | 2.48 | 0.013 |
| geniculate_thalamus | L | Thalamus | Subcortical | 2.48 | 0.013 |
| area_TEm | R | Temporal | Limbic | 2.47 | 0.013 |

[1] Sallet, J., Mars, R. B., Noonan, M. P., Andersson, J. L., O'Reilly, J. X., Jbabdi, S., Croxson, P. L., Jenkinson, M., Miller, K. L., and Rushworth, M. F. S. (2011). Social network size affects neural circuits in macaques. *Science*, 334(6056):697–700.

[2] Testard, C., Brent, L. J. N., Andersson, J., Chiou, K. L., Negron-Del Valle, J. E., DeCasien, A. R., Acevedo-Ithier, A., Stock, M. K., Antón, S. C., Gonzalez, O., Walker, C. S., Foxley, S., Compo, N. R., Bauman, S., Ruiz-Lambides, A. V., Martinez, M. I., Skene, J. H. P., Horvath, J. E., Unit, C. B. R., Higham, J. P., Miller, K. L., Snyder-Mackler, N., Montague, M. J., Platt, M. L., and Sallet, J. (2022). Social connections predict brain structure in a multidimensional free-ranging primate society. *Sci. Adv.*, 8(15):eabl5794.
